# Genome-wide mapping of DNA G-quadruplexes in *Trypanosoma brucei* chromatin reveals enrichment in coding regions

**DOI:** 10.1101/2025.07.22.666098

**Authors:** Ludovica Monti, Genta Firth, Joana R. Correia Faria, John M. Kelly, Gem Flint, Silvia Galli, Thomas E. Maher, Marco Di Antonio

## Abstract

G-quadruplexes (G4s) are non-canonical DNA structures formed in guanine-rich sequences that are proposed to act as regulatory elements in trypanosomatid parasites, including *Trypanosoma brucei*, the causative agent of African sleeping sickness. However, their functional roles remain poorly understood, largely due to limited knowledge of their genomic distribution. Herein, we performed computational analyses across 63 trypanosomatid species uncovering high degree of variability in G4-prevalence and species-specific patterns. We generated the first genome-wide map of G4s in *T. brucei* using G4 chromatin immunoprecipitation followed by sequencing (G4 ChIP-Seq), which revealed a striking enrichment of G4s within coding DNA sequences (CDSs). This pattern diverges markedly from *in silico* predictions and previous genome-wide G4 mapping studies in humans, suggesting that G4s may play unique roles exclusive to trypanosome biology. To investigate their functional relevance, we profiled the transcriptome of *T. brucei* upon treatment with the G4-stabilising ligand PhenDC3. We observed that PhenDC3 exerts targeted gene expression perturbation of genes bearing G4s, particularly those located within coding CDSs, where G4s are mostly enriched. Altogether, our findings highlight a distinctive role for G4s in the regulation of gene expression in *T. brucei* and support their potential as therapeutic targets in the treatment of African sleeping sickness.

**GRAPHICAL ABSTRACT:** 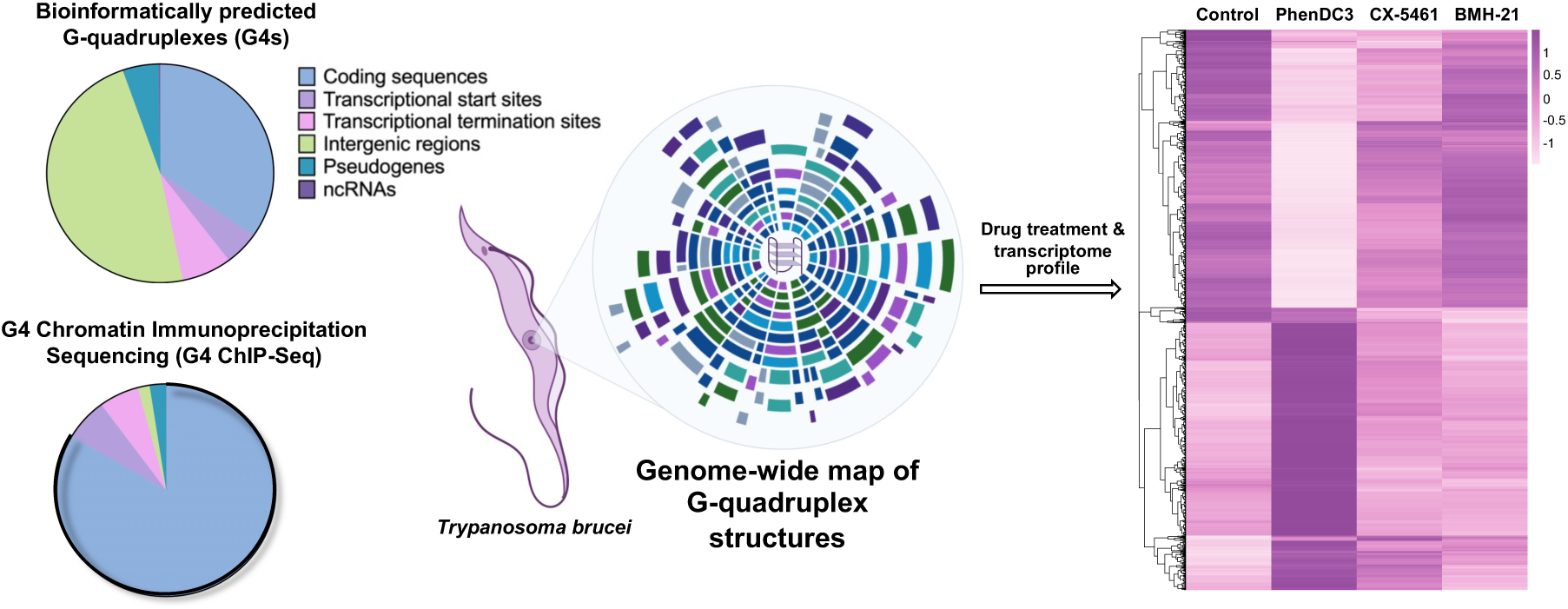

## INTRODUCTION

Human African trypanosomiasis, commonly known as "sleeping sickness," is a parasitic disease caused by protozoa of the *Trypanosoma* genus, predominantly transmitted through the bite of infected tsetse flies. This disease is classified as one of the neglected tropical diseases (NTDs), significantly affecting impoverished populations in sub-Saharan Africa [1,2]. An estimated 55 million people in 36 sub-Saharan African countries are at risk, with approximately 1,000 cases reported annually [1]. The disease often progresses to late-stage infections that affect the central nervous system causing neurological damage and is fatal without timely treatment [3]. Current treatments for African trypanosomiasis present significant challenges, including toxicity, complex administration, and the need for hospitalization [4,5]. The emergence of drug-resistant strains poses further risks, especially in rural areas where access to healthcare is limited [6-8]. Fexinidazole, introduced in 2019 as an oral treatment marked a significant advancement in simplifying treatment protocols [9]. However, its efficacy in advanced-stage disease, as well as in children below a certain age and weight, pregnant women, and breastfeeding mothers remains unclear [10,11].

The impact of African trypanosomiasis extends beyond human health, affecting agriculture and economic stability across the region [12]. Animal trypanosomiasis, known as Nagana, has severe consequences for the livestock industry, particularly cattle, where weight loss, reduced productivity, and high mortality, leads to economic losses in agriculture exceeding $1 billion annually [13,14]. These growing concerns highlight the urgent need for innovative therapeutic strategies. Identifying new molecular targets is therefore essential to addressing these challenges and advancing the development of safer, more effective treatments against drug-resistant strains.

G-quadruplexes (G4s) are DNA secondary structures formed in guanine-rich sequences through self-assembly mediated by Hoogsteen hydrogen bonding [15]. These structures consist of stacked guanine tetrads, which are stabilised by the presence of central monovalent cations such as K^+^ or Na^+^[16,17]. G4 formation is typically observed in genomic sequences containing at least three guanine tetrads separated by loops of up to seven nucleotides (GGG-N_1-7_-GGG-N_1-7_-GGG-N_1-7_) [18]. However, recent experimental evidence has revealed that sequences deviating from this general consensus motif can also form stable G4s under physiological conditions, which are often referred as non-canonical G4s [19]. G4s are non-randomly distributed across the genomes of all eukaryotic organisms. In humans, predictive algorithms [20] and next-generation sequencing techniques [19,21] have revealed enrichment of G4s in regulatory regions, such as telomeres and promoters of proto-oncogenes (i.e., *c-MYC* [22] and *c-KIT* [23]). These findings underscore the potential relevance of G4s in key cellular processes, including transcriptional regulation, replication, and genome stability [17,24]. The development of genome-wide mapping strategies to underpin the distribution of G4s in chromatin have further revealed how the formation of these structures has a direct impact in promoting transcription, especially when formed at promoters [25,26]. More recently, we identified that clusters of G4s formed at intergenic and intronic regions can act as super enhancers (super-G4s), further highlighting a potential role of G4s as epigenetic marker for transcriptional control [27]. Based on these observations, significant efforts have been made in the development of G4-targeting drugs [28,29], with two candidates – CX-5461 and QN-302 – progressing through clinical trials for the treatment of tumour malignancies [30-32], underscoring the translational impact of G4-targeting therapeutic approaches. This has promoted the investigation of G4-targeting beyond cancer treatment and across various organisms, including pathogens (e.g., viruses, bacteria, parasites) [33-35], and others (e.g., yeasts, helminths, plants) [21,36,37]. For example, in protozoan parasites such as *Plasmodium falciparum* (the causative agent of malaria) and *T. brucei*, putative G4s have been identified in the surrounding regions of virulence genes and trypanosomatid-specific epigenetic modifications (i.e., *base J*) linked to parasite antigenic variation and survival in their hosts [33,37-39]. This has prompted the development of a series of new G4-ligands with anti-proliferative effects in *T. brucei* cells, highlighting the potential of G4s as therapeutic targets against *Trypanosoma* infections [40-50]. Despite these promising findings, underpinning the molecular mechanism that justify the effectiveness of G4-ligands to treat *Trypanosoma* infections remains challenging due to the absence of a reliable genomic map of G4 distribution in this parasite. This limits our ability to fully harness the therapeutic potential of these DNA structures for treating sleeping sickness and other trypanosomatid infections.

In this study, we present the first genome-wide mapping of G-quadruplex (G4) structures in *T. brucei* chromatin and explore their functional relevance. Through G4-targeting chromatin immunoprecipitation followed by sequencing (G4 ChIP-Seq) and transcriptomic profiling, we reveal that G4s are widely distributed across the *T. brucei* genome, with a notable enrichment in coding sequences (CDSs)—a localisation that diverges from the promoter-centric distribution seen in mammals. Functional assays using the G4-stabilising ligand PhenDC3 showed selective upregulation of G4-containing genes, particularly within CDSs, suggesting a distinct mechanism of G4-mediated gene regulation in *T. brucei*. These findings underscore the potential of targeting G4 structures to perturb essential gene networks in trypanosomes and inform novel strategies for therapeutic intervention.

## MATERIAL AND METHODS

### *In silico* bioinformatic analyses

*Genome sequences.* A total of 63 genomes representing seven trypanosomatid genera were downloaded from the NCBI genome database: *Trypanosoma*, *Leishmania*, *Crithidia*, *Phytomonas*, *Herpetomonas*, *Leptomonas*, and *Porcisia*. Additionally, three non-trypanosomatid genomes were included for comparative analyses: two *Schistosoma* genomes [51] (a trematode parasite that causes a neglected tropical disease) and the human genome [52] for comparison. Accession numbers for all genomes are provided in Supplementary Table S1.

*G4Hunter analysis.* The G4Hunter algorithm [53-55] was used to identify putative G-quadruplex sequences (PQSs) across multiple genomes. A custom Python script was developed to automate the predictive analysis in iterative loops. The analysis utilized two key input parameters: window size and threshold. The window size defines the length of DNA fragments analysed, while the threshold determines the propensity of a sequence to form a G4 structure. Unless otherwise specified, the default G4Hunter settings of 25-nucleotide window size and a threshold of 1.2 were used, as these parameters have been previously validated to achieve a balance between minimizing false positives and false negatives [51,53]. To further refine the analysis, additional evaluations were conducted at higher thresholds of 1.5, 1.8, and 2.0. The frequency of PQSs per 1000 base pairs for each species was determined by diving the number of PQSs identified using G4Hunter by the corresponding genome length, and multiplying the result by 1000.

### Cell cultures

#### Trypanosoma brucei

*T. brucei* Lister 427 bloodstream forms were grown in HMI-11 medium (Gibco) supplemented with 10% fetal bovine serum (FBS, Sigma-Aldrich) at 37°C and 5% CO_2_. The density of cell cultures was maintained below 1.5 × 10^6^ cells/mL and sub-passaged every 24–48 hours.

#### HeLa cells

HeLa cells were grown in culture flasks or microtiter plates after initial seeding at 1 × 10^4^ cells/mL in DMEM medium (Sigma-Aldrich) with 10% FBS. They were sub-cultured every 5 days.

### Preparation of recombinant BG4 antibody

Recombinant BG4 was expressed as previously described [56]. Briefly, BL21(DE3) *E. coli* cells were inoculated with the expression vector pSANG10-3FBG4 (Addgene, plasmid no. 55756) [57]. via heat shock, then grown overnight in 2xTY medium containing glucose (2%) and kanamycin (50 mg/ml, Gibco) before expanding into autoinduction medium. The culture was grown at 37°C for 6 hours, followed by overnight incubation at 25°C. After harvesting the cells, protein purification was performed using Nickel Affinity Chromatography, elution in the presence of imidazole, followed by dialysis in PBS overnight. Protein purity and concentration were assessed by SDS-PAGE, and antibody functionality was evaluated by ELISA [58].

### G4-targeting chromatin immunoprecipitation followed by sequencing (G4 ChIP-Seq)

Cell lysis and chromatin fragmentation were carried out as in ref. [59,60]. Briefly, 2 × 10^8^ parasites were cross-linked with 1% formaldehyde for 20 minutes at room temperature. Chromatin was sonicated using a Bioruptor (Diagenode) with sonication beads (Diagenode) for 20 cycles (30 seconds on, 30 seconds off) using the high setting. Cell debris were pelleted by centrifugation, and 10% TritonX was added to each sample. Assessment of successful chromatin sonication to a required fragment size distribution of 100-500 bp was carried out using either a 2% (wt/vol) agarose gel electrophoresis or the Agilent 4200 TapeStation system (version 4.1.1) and the Agilent D5000 screentape following manufacturer’s instructions. Aliquots of chromatin were flash frozen in liquid nitrogen and stored at -80 °C until further use.

Chromatin immunoprecipitation (ChIP) was carried out as in ref. [58] with the following modifications. Chromatin (76.2 ng/μL) was thawed on ice before being supplemented with Triton X-100 (2%) and incubated at room temperature for 10 minutes. Chromatin (9.4 μL/sample; 720 ng/sample), RNase A (1 μL/sample, Ambion^TM^) and blocking buffer (37 μL/sample; containing: intracellular salt buffer with 1% Bovine Serum Albumin (BSA) in Milli-Q water, filter sterilised before use) was incubated on a thermomixer under rotation (1,400 r.p.m.) at 37°C for 20 minutes. BG4 (0.75 μL/sample; 136 ng/sample) was added to the ChIP samples. One sample was left untreated with BG4 and kept on ice to serve as the input (IP) control sample. The ChIP samples treated with BG4 were incubated at 16°C with 1,400 r.p.m. rotation for 1 hour. Meanwhile, Anti-FLAG antibodies (5 μL/sample, Millipore) were washed three times with blocking buffer (50 μL/sample) using the DynaMag-2 magnet and then resuspended in blocking buffer (50 μL/sample). Anti-FLAG bead suspension (50 μL/sample) was added to the ChIP samples and incubated at 16°C with 1,400 r.p.m. rotation for 1 hour. The ChIP samples were then washed three times at 4°C with ice-cold wash buffer (200 μL/sample; containing: 100 mM KCl, 0.1% (vol/vol) TWEEN-20 and 10 mM Tris, pH 7.4, in Milli-Q water, and filter sterilised before use). Two additional washes were performed using wash buffer (200 μL/sample), followed by a 10-minute incubation at 37°C and 1,400 r.p.m. TE buffer was then added to all samples (including the input) to a final volume of 75 μL, followed by the addition of 1 μL of Proteinase K (20 mg/mL, Invitrogen). The samples were incubated for 1 hour at 37°C with continuous rotation at 1,400 r.p.m., followed by a 2-hour incubation at 65°C with 1,400 r.p.m. rotation. Samples were then purified using MinElute Reaction Cleanup Kit, following the manufacturer’s instructions, with elution in 25 μL of elution buffer. The DNA concentration of the samples was measured using a Qubit fluorometer with the Qubit dsDNA HS Kit (Thermo Fisher Scientific, MA, USA). Using chromatin derived from a single *T. brucei* culture (representing one biological sample), three technical replicates (ChIP samples) and one IP sample were performed in parallel.

Prior to sequencing, quantitative PCR (qPCR) was used to determine enrichment of G4-forming regions over non-G4-forming regions in ChIP samples compared to IP sample. Purified DNA samples were diluted in DNase/RNase-free water, and 5 μL of the diluted DNA was amplified via qPCR using 10 μL of Fast SYBR Green Master Mix and 2.5 μL each of forward and reverse primers at a final concentration of 1 μM (primer sequences listed in Table 1). Technical duplicates were tested for each primer set in both ChIP and IP samples. qPCR was performed on a Stratagene Mx3005P Real-Time PCR system (Agilent Technologies, CA, USA) using the following program of initial denaturing at 95°C for 20 seconds, followed by 40 cycles of denaturing at 95°C for 3 seconds, and annealing and extension at 60°C for 30 seconds.

**Table 1.**
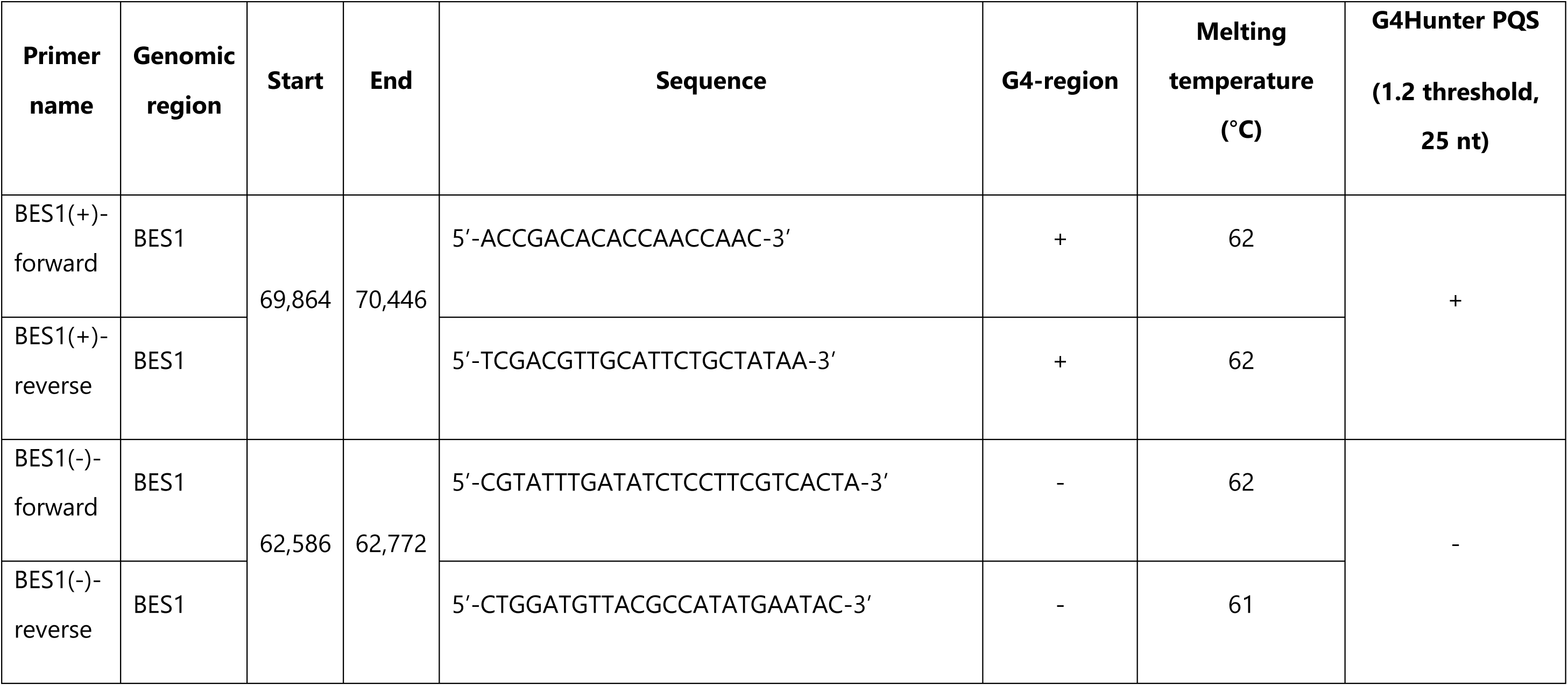
Forward and reverse primers used for quantitative PCR (qPCR) to assess G-quadruplex (G4) enrichment at G4-positive and G4-negative loci within the bloodstream expression site 1 (BES1) of *T. brucei*.

To calculate the fraction of recovery (% Input), Ct values were adjusted relative to the input sample using the formula: 2^(Ct^ ^value^ ^of^ ^the^ ^Input–Ct^ ^value^ ^of^ ^the^ ^ChIP)^ x 100. To estimate the G4-fold enrichment, the % Input of a G4-positive region was divided by the % Input of a G4-negative region. Fold enrichment was then calculated by dividing the average percentage input of G4 regions by that of the non-G4 regions.

Libraries were prepared from input and immunoprecipitated DNA using the NEBNext Ultra II DNA Library Prep Kit for Illumina (New England Biolabs, MA, USA). Final libraries were sequenced on the Illumina NovaSeq 6000 (S2 200 cycles paired end reads) at a depth of 15 million reads per sample.

#### ChIP-Seq data analysis

Recombinant Read quality was assessed using *FastQC* (http://www.bioinformatics.babraham.ac.uk/projects/fastqc/), and adapter sequences were trimmed with *fastp* [61]. Trimmed reads were aligned to the 2018 genome assembly of the Lister 427 version 10 [62]. (Tb427v10) extracted from TriTrypDB, using *Bowtie2* [63] with default settings. Reads were filtered and sorted using *SAMtools* [64] with the following arguments *-F 2308* and *-q 10* to exclude unaligned, secondary, and supplementary reads, and reads with a mapping quality (MAPQ) < 10. Duplicate reads were removed using *Picard MarkDuplicates* (https://broadinstitute.github.io/picard/) with the *REMOVE_DUPLICATES=TRUE* option. *Bowtie2* alignment files were analysed for peak calling (i.e., read enrichment) using *MACS2* [65] with a user-defined genome size of -g 2.67e7; 200 nt fragment size for shifting model (--nomodel --extsize 200); minimum false discovery rate: ≥0.01 (-q 0.01). Log_2_ ratio of ChIP signal to Input signal were generated using *bamCompare* from the *deepTools* [66] suite (version 2.4.2) with a bin size of 50 bp. The resulting bigWig files were visualised using the Integrative Genomics Viewer (IGV, version 2.19.4) [67]. *Bowtie2* and *MACS2* output files were also visualised with IGV. Common and unique peaks between replicates were identified with *bedtools intersect* [68].

#### Genomic feature annotation and enrichment analysis

Genomic distribution of ChIP-Seq peaks was annotated using the *GenomicRanges* package in R [69]. Peaks were assigned to genomic features based on the extent of their overlap with annotated regions. Gene feature annotations were based on the *T. brucei* 427 genome assembly Tb427v10 [62]. To supplement this, 341 of the 342 annotated transcription start sites (TSS) and transcription termination sites (TTS) from the latest genome assembly (v12) [70] were successfully mapped to the v10 assembly using *Liftoff* [71]. Intergenic regions were defined as genomic intervals not overlapping any annotated gene feature (i.e., protein coding genes, TSS, TTS, pseudogenes, ncRNA) and were included as a separate feature category in the annotation and enrichment analysis.

To assess genomic enrichment, random genome-wide shuffling of ChIP-Seq G4-peaks was performed three independent times using *bedtools shuffle* (v2.31.1) [68], and the overlap was assessed in each randomised case. Fold enrichment was calculated for each genomic feature by dividing the observed overlap of ChIP-Seq peaks by the overlap from each of three randomized datasets, producing three-fold enrichment values. The standard deviation of these values was then computed. For polycistronic transcription unit (PTU) analysis, gene-to-PTU associations were constructed using genome annotations from Tb427v12. A gene–PTU mapping dictionary was generated in R using the *GenomicRanges* package [69], allowing each gene to be linked to its corresponding PTU for downstream enrichment and localization analysis.

Radial plots for the ChIP-Seq data were generated using *Circos* hosted on Galaxy [72,73], and edited using *Affinity Designer 2* (https://affinity.serif.com). The genomic regions were based on coordinates defined by the official genome FASTA assembly, excluding unitig and unplaced-contig fragments. Bloodstream expression sites (BESs) were treated as a distinct track due to their specialised and independently regulated nature within the subtelomeric domains. The log_2_ fold change (ChIP/Input) across the genome was calculated using *bamCompare* from the *deepTools* [66] suite with a bin size of 1000 bp, generating bigWig files for each ChIP replicate. Replicate bigWig files were then averaged using *bigwigAverage* function from *deepTools* [66], and the resulting mean bigWig file was converted to WIG format using *bigWigToWig* via Galaxy [74].

### Cell viability assays with G4-ligands

Antitrypanosomal activity against *T. brucei* Lister 427 bloodstream forms, and cytotoxicity in HeLa cells were assessed for selected G4-targeting molecules. BMH-21 and CX-5461 were purchased from Cayman Chemical (Item No. 22282, MI, USA) and Selleckchem.com (Item No. S2684, TX, USA), respectively. PhenDC3 was synthesised in the lab following established procedures [75]. The viability of *T. brucei* and HeLa cells were evaluated in 96-well microtiter plates (200 μL per well). Each viability assay was initiated with 2.5 × 10^4^ parasites/mL or 1 × 10^4^ HeLa cells/mL, followed by the addition of test compounds at concentrations ranging from nM to μM. For *T. brucei,* plates were incubated for 48 hours, after which resazurin (20 μL at 0.125 mg/mL) was added, and the plates were incubated for an additional 16 hours. For HeLa cells, plates were incubated for 5 days, resazurin was added (as above), and the plates incubated for another 6 hours. Fluorescence was measured using a BMG FLUOstar Omega plate reader (excitation at 545 nm, emission at 590 nm). EC_50_ and CC_50_ values were calculated using GraphPad Prism and are shown as the mean ± standard deviation (SD). Assays were performed in triplicates.

### RNA-sequencing

*T. brucei* Lister 427 bloodstream forms (0.75 × 10^6^/ml; 15 ml) were treated with 2 × EC_50_ concentrations of either PhenDC3, CX-5461, BMH-21, or without compound (control) for 4 hours. Four replicates were prepared for each experimental condition. Total RNA was extracted using the RNeasy Mini Kit (Qiagen, Australia), following the manufacturer’s protocol. Briefly, cell pellets were lysed with RNeasy lysis buffer and homogenized using QIAshredder columns. The lysate was subsequently washed with RNeasy wash buffer, and on-column DNase I digestion was performed to remove genomic DNA. RNA was purified and eluted in 50 μL of RNase-free water. RNA quality was assessed using a DeNovix spectrophotometer. Poly(A)-enriched RNA libraries were prepared and sequenced on the Illumina NovaSeq 6000 (S2 200 cycles paired end reads) at a depth of 15 million reads per sample.

#### RNA-seq data analysis

The quality of sequence reads was assessed using *FastQC* (http://www.bioinformatics.babraham.ac.uk/projects/fastqc/), and sequence files were concatenated using the *cat* function (Unix version 8.30). Sequencing adapter sequences were trimmed using *fastp* [61]. Trimmed reads were aligned to the 2018 genome assembly of the Lister 427 strain version 10 (Tb427v10, https://tritrypdb.org) using *Bowtie2* [63]. The alignments were converted from SAM to BAM format, sorted, indexed and filtered by alignment quality ≥ 1 (-q 1) using *SAMtools* [64]. Reads were quantified using f*eatureCounts* [76] from the *subread* package (v1.5.2). Differential gene expression analysis and volcano plot visualisations were performed using *edgeR* package in R [77], applying a False Discovery Rate (FDR) threshold of 0.05 and a log_2_ Fold Change (FC) cut-off of ±1 (corresponding to a minimum twofold change in expression). Volcano plots were generated using the *ggplot2* [78] package in R. For the analysis of key genes, gene expression data for each experimental condition were exported as CSV files, and intersected with ChIP-Seq data. GraphPad Prism software was used to visualise the downstream statistical analyses. Multi-Dimensional Scaling (MDS) plot was generated with *Degust* [79] using min gene read count of 10, FDR threshold of 0.05 and abs log FC cut-off of 1.

### Data deposition

ChIP-Seq and RNA-seq data have been deposited at gene expression omnibus (GEO), under accession number GSEXXXXXX.

## RESULTS

### G-quadruplexes are abundant in all trypanosomatids

While current studies often focus on a limited set of representative genomes from trypanosomatids, we have extended the scope of predictive analyses by examining the frequency of putative G-quadruplex sequences (PQSs) per kilobase across the genomes of seven trypanosomatid genera (*Trypanosoma*, *Leishmania*, *Crithidia*, *Phytomonas*, *Herpetomonas*, *Leptomonas*, and *Porcisia*) using the G4Hunter algorithm [53-55]. This comprehensive approach allows for a broader investigation of G4 distribution within the trypanosomatid family and can be leveraged to highlight changes in their prevalence across different species. For comparative analyses, we also included two *Schistosoma* [51] genomes and the *Homo sapiens* [52] genome, as representatives of other parasitic and mammalian organisms. The length of trypanosomatid genomes selected for the analyses varied from approximately 17 Mb (*Phytomonas francai*, smallest) to 89 Mb (*T. cruzi*, biggest), as shown in Supplementary Table S1. In contrast, the genome of schistosomes and humans is approximately 4–35 times larger (400 Mb, and 3.1 Gb, respectively). As displayed in Figure 1A, we observed high variability in PQS prevalence across genera, with *Herpetomonas* and *Leishmania* species exhibiting the highest G4-frequencies and *Schistosoma* showing comparatively the lowest G4-prevalence. Specifically, the highest frequency of G4s (as number of PQSs per kbp) was found in *Herpetomonas tarakana*, which exhibited a 27-fold higher abundance of G4s (frequency of 5.63) compared to *Schistosoma japonicum* (frequency of 0.22) and a six-fold higher abundance than in humans (frequency of 0.93, Figure 1A). The PQS abundance observed in trypanosomatids compared to schistosomes and humans could, in part, be explained by their higher GC content (average GC% = 52.9%) relative to *Schistosoma* (average GC% = 32.9%) and humans (GC% = 37.8), and therefore a higher propensity to form G4 structures. Visualising the GC content against the frequency of PQSs revealed distinct patterns across genera (Figure 1B), showing a near linear relationship between GC richness of a given genome and its G4 prevalence (R squared value of 0.709). Interestingly, *Leishmania* species exhibited little variation in G4 prevalence across all evaluated species (Figure 1B), suggesting potential evolutionary conservation of G4s within this genus that may reflect their relevance to the biology of the parasite. In contrast, *Trypanosoma* species showed greater variability in G4 frequencies (inset zoom in Figure 1B), suggesting potential differences in the biological and therapeutic significance of G4s across these species. The variation in G4 density observed within the *Trypanosoma* genus suggests that G4 prevalence does not always correlate with the GC richness of these parasites, supporting the notion of a potential need of these structures for the biology of certain species. For example, *T. cruzi* and *T. rangeli* have comparable GC content (51.55% and 51.52%, respectively), whilst displaying a 1.3-fold difference in G4 prevalence (*i.e.,* 1.96 and 2.60 PQS frequency per kbp, respectively). The variability in G4 distribution detected suggests that the GC content alone cannot be used to reliably predict the abundance of G4s in each species, which might be determined by other factors. For example, while both species can infect humans, *T. rangeli* is non-pathogenic to mammals [80], suggesting that variations in G4 abundance may be linked to differences in their biological functions, potentially regulating pathogenic potential or disease-causing mechanisms.

**Figure 1:**
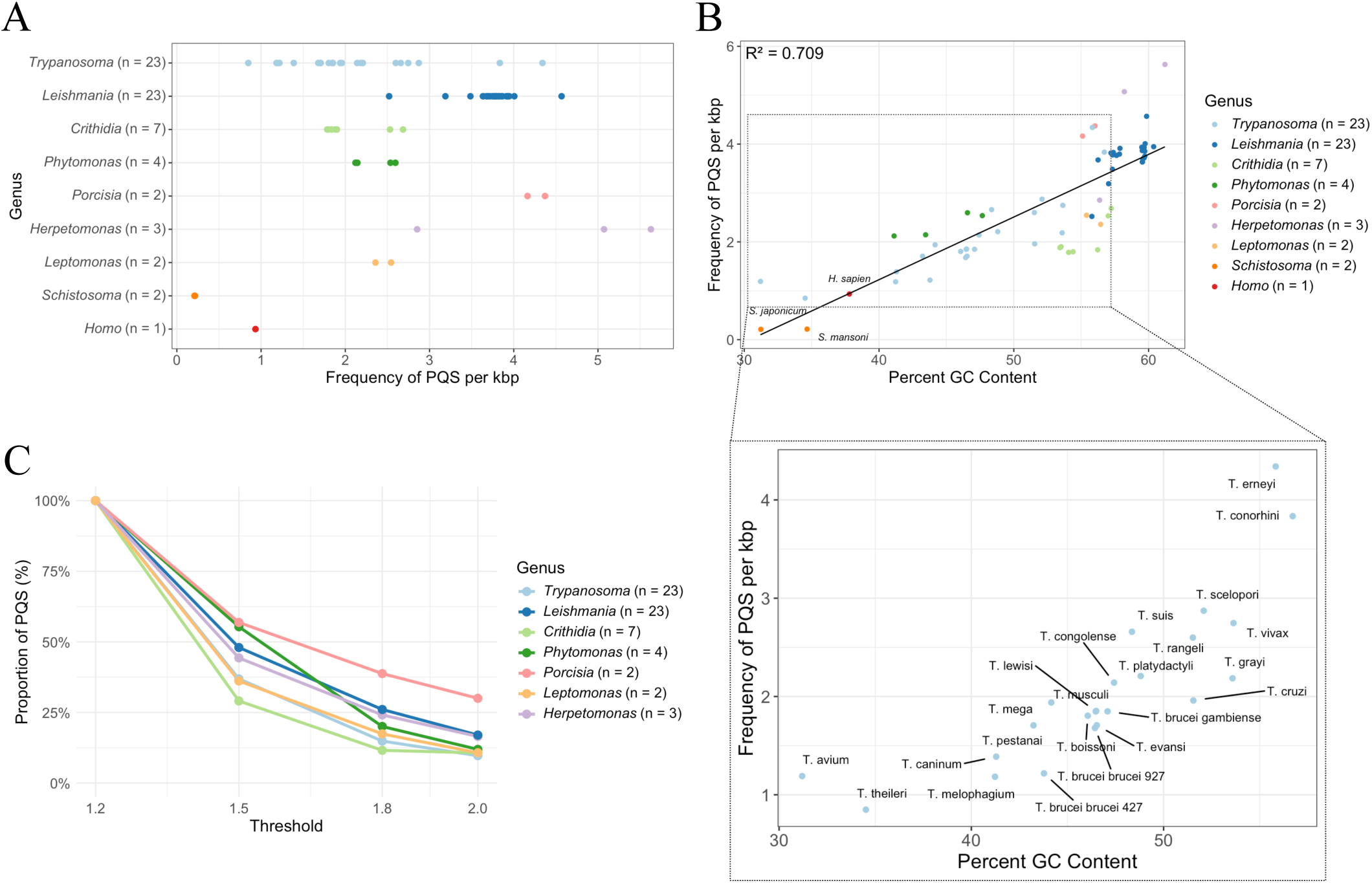
Whole-genome *in silico* analysis of putative G4-forming structures across trypanosomatid species. A) Frequency of PQSs per kilobase (kbp). Each dot represents individual species genomes within the analysed trypanosomatid genera, as specified on the Y axis. The number of species analysed per genus is shown in parentheses. For comparison, *Schistosoma* and *H. sapiens* genomes are included as reference controls. B) Frequency of PQSs plotted against GC content of each trypanosomatid species analysed. The main graph shows results for all species, while the inset zooms in on *Trypanosoma* species, highlighting the variability in PQS frequency and GC content. C) Change in the proportion of PQSs retained at increasing G4Hunter thresholds (1.2, 1.5, 1.8, and 2.0), expressed as a percentage (%) relative to the mean number of PQSs identified at the lowest threshold (1.2). For each trypanosomatid genus, the mean PQS count at each threshold was divided by the mean count at threshold 1.2 and multiplied by 100, highlighting the sensitivity of PQS detection to threshold stringency.

To assess the propensity of DNA sequences to form G4 structures, we performed additional analyses by adjusting the threshold values in G4Hunter, which quantifies the likelihood of a DNA sequence to form a G4 structure based on its guanine content and arrangement [53]. Higher G4Hunter threshold values correspond to sequences with a stronger likelihood of forming stable G4 structures; conversely, a lower threshold value predicts G4 structures that are, in general, less stable [51,53]. As shown in Figure 1C, increasing G4Hunter thresholds from 1.2 to 2.0 resulted in a progressive decrease in the sensitivity of the analysis (i.e., proportion of PQS found). *Porcisia* retains the largest proportion of PQSs across all thresholds, demonstrating high genomic density of sequences capable of forming stable G4s. It is worth noting that *Porcisia* does not have the highest GC content among the species analysed, which further suggests that GC richness alone is not the primary determinant of G4 abundance, and that evolutionary enrichment of G4s in these parasites might reflect their biological significance.

### *T. brucei* G-quadruplexes (G4s) are enriched in coding regions

Given the similar PQS abundance in human and *Trypanosoma* cells, and the variability of G4 prevalence among the 22 *Trypanosoma* species examined–despite their comparable GC content–we decided to experimentally map the genomic distribution of G4s in *T. brucei* to gain further insight into their biological roles and potential contribution to pathogenicity. To achieve this, we adapted an established G4-targeting Chromatin Immuno-Precipitation sequencing (G4 ChIP-Seq) protocol [58] to enable the first detection of G4s within the chromatin of bloodstream forms of *T. brucei* parasites. Briefly, chromatin was extracted from *Trypanosoma* cells following an established protocol [59,60] and crosslinked with formaldehyde to preserve its native architecture. The concentration of recombinant G4-selective antibody, BG4 [57], was optimised empirically to enhance the enrichment of G4-containing chromatin fragments while minimising non-specific binding. Key refinements to the G4 ChIP-Seq protocol established for human chromatin [58] included: adjustments to chromatin-to-antibody ratios (∼ 5:1), chromatin pre-treatment with Triton X-100 (2% for 10 min), and optimisation of sonication cycles (i.e., 20 cycles at 30 seconds on and 30 seconds off) to achieve optimal chromatin fragmentation of 100–500 bp. Fine-tuning of these parameters was essential to successfully perform G4 ChIP-Seq on nanograms of trypanosome chromatin, which is imposed by the low *T. brucei* culture yields. Next, we assessed G4 ChIP quality by measuring the relative enrichment of predicted G4-positive *versus* G4-negative regions, which were identified through a pilot ChIP-Seq study (see Materials and Methods). These regions were located within the bloodstream expression site 1 (BES1), a telomeric locus in *T. brucei* that controls the expression of the active variant surface glycoprotein (VSG) gene, a key factor in immune evasion [62,81]. Using our adapted G4 ChIP protocol for *T. brucei*, we observed approximately 6-fold enrichment of G4 containing regions relative to non-G4 controls, consistent with the ≥5-fold enrichment benchmark established for successful G4 immunoprecipitation in human cells (Figure 2A) [58]. DNA libraries were then prepared and sequenced using Illumina technology at a depth of 15 million reads per sample.

**Figure 2:**
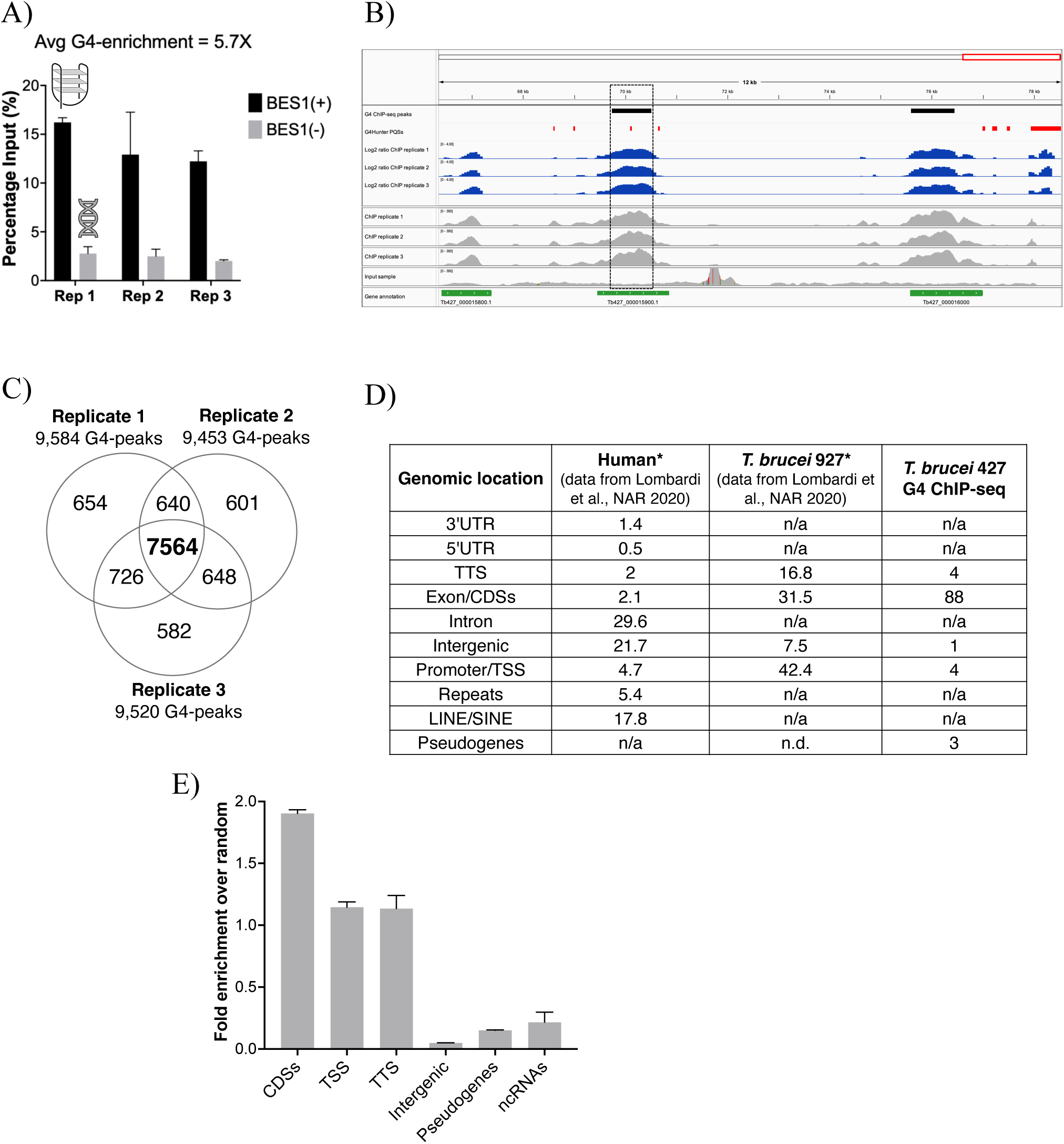
G4-targeting Chromatin Immuno-Precipitation followed by sequencing (G4 ChIP-Seq) in trypanosomes. A) G4 enrichment in *T. brucei* chromatin assessed by qPCR using primers targeting G4-positive (BES1(+)) and G4-negative (BES1(-)) regions. The average G4 enrichment across three ChIP technical replicates is 5.7-fold relative to control sequences lacking G4 structures. B) Integrative Genomics Viewer snapshot of the bloodstream expression site 1 (BES1) region (66,342-78,519), showing enrichment of G4-associated peaks identified by ChIP-Seq (in black) relative to the reference input control. Peaks were detected using the peak-calling algorithm *MACS2* [65]. Overlapping putative G4-forming sequences (PQSs, in red) were predicted using G4Hunter with a stringency threshold of 1.2 and a 25 nucleotide window. Log_2_(ChIP/Input) signal ratios, generated using *bamCompare* from the *deepTools (66)* suite, are shown in blue; while ChIP and input BAM file coverages (indexed using *samtools*) are displayed in grey. The dashed box highlights the G4-positive *locus* targeted by primers for qPCR validation of G4 enrichment post-ChIP. Genome functional annotations are shown in green. C) Venn diagram illustrating the number of G4-forming sequences (i.e., G4-peaks) identified in *T. brucei* through G4 ChIP-Seq across three technical replicates. The three replicates show a high degree of overlap, with 79% of G4-peaks (7,564 peaks) shared among them, highlighting the high reproducibility of the technique. D) Table illustrating the genomic distribution of G4s, as identified through previous experimental and *in silico* analyses in human and *T. brucei* (data from ref. [52]), compared to G4-regions identified through ChIP-Seq. E) Bar plot showing fold enrichment of ChIP-Seq G4-peaks in different genomic regions, such as coding DNA sequences (CDSs), transcriptional start sites (TSS), transcriptional termination sites (TTS), intergenic regions, pseudogenes, and non-coding RNAs (ncRNAs). Mean and Standard Deviation (SD) values are shown.

Sequencing data were aligned to the *T. brucei* reference genome (2018 version 10), using *SAMtools* [64] and applying a high mapping quality (MAPQ) threshold of 10 to prioritise uniquely mapped reads. G4-enriched regions were identified with the peak-calling program *MACS2* [65], using parameters [82] optimised for the *T. brucei* genome: size was set to 2.67 × 10^7^, a stringent q-value threshold of ≤0.01 (-q 0.01) was used to minimise false positives, and a fixed fragment size of 200 nucleotides (--extsize 200) was applied. This analysis allowed for the detection of G4-peaks (Figure 2B), providing the first comprehensive map of G4s in *T. brucei* chromatin. Our workflow yielded consistent and reproducible G4-enrichment patterns across three independent ChIP replicates (Figure 2 B).

Experimentally derived G4 data were compared with previously published *in silico* predictions and genomic maps of G4s in human and *T. brucei* genomes to assess differences in their distribution patterns [21,52]. Our G4 ChIP-Seq analysis identified an average of 9,519 peaks in *T. brucei* chromatin across three technical replicates (Figure 2C), a number substantially lower than the ∼30,000 PQSs predicted by computational analysis [52]. High reproducibility was observed, with 7,564 G4 peaks (79%) consistently shared across all three replicates (Figure 2C). Moreover, 36% of the experimentally identified G4-peaks overlapped with PQSs predicted *in silico* by G4Hunter (Supplementary Table S2). We then compared the genomic distribution of G4s revealed by our ChIP-Seq data to patterns previously reported previously in studies using *in silico* and *in vitro* approaches. G4-seq performed by Marsico *et al.* [21] on naked single-stranded DNA fragments of the *T. brucei* genome revealed a similar G4 distribution pattern between the human and the parasite genomes. In a subsequent *in silico* study, Puig Lombardi *et al.* [52] intersected the G4 sequences identified by G4-seq [21] with those predicted by G4Hunter, and annotated their genomic distribution. This study found a high prevalence of G4s in promoters (43%) and exons (32%) of the *T. brucei* genome, compared to the high enrichment in regulatory regions, such as promoters and introns, observed in humans (Figure 2D) [52]. Although previous studies provided valuable insights into the distribution of G4s in *T. brucei* and humans, direct comparison of genomic locations is challenging due to the unique genomic organisation of the parasites. The parasite’s genes are organised in long polycistronic transcription units (PTUs), containing dozens of functionally unrelated genes. Unlike many eukaryotes, *T. brucei* lacks classical promoters for individual genes; instead, transcription initiates at defined transcription start sites (TSS) located upstream of these PTUs [83]. Furthermore, genome annotation uses coding DNA sequences (CDSs) rather than exons, as *T. brucei* does not possess the typical exon-intron gene structures common in other eukaryotes. This distinct genomic architecture requires a more rigorous and tailored approach to accurately locate G4s within the parasite’s genome based on the ChIP-Seq data. To enhance annotation accuracy, we employed a combined approach whereby ChIP-Seq peaks were initially annotated using the *T. brucei* Lister 427 genome 2018 assembly [62] (version 10, extracted from TriTrypDB) as reference, which includes protein coding regions, pseudogenes, and non-coding RNAs (ncRNAs).

Subsequently, transcription start sites (TSS) and transcription termination sites (TTS) from the most recent genome assembly (v12) [70] were mapped onto the v10 assembly using *Liftoff* [71]. Lastly, peaks that did not overlap any of the previously annotated gene features were classified as intergenic regions. Our analysis revealed a markedly different pattern of G4 distribution in the *T. brucei* genome compared to previous studies, suggesting that G4 formation in the living parasite might be driven by biological processes regulated by these structures. Notably, our data indicate that G4s are predominant within CDSs, accounting for 88% of detected peaks. This represents a substantially higher proportion than the 32% and the 2.1% reported by Puig Lombardi *et al.* in *T. brucei* and humans, respectively (Figure 2D) [52]. We also observed a considerably lower number of G4s located at TSS and TTS, with only 4% of peaks localising to these regions each. This again contrasts with previous studies by Puig Lombardi *et al.*, which reported 42% of potential G4-sequences at TSS and 17% at TTS in *T. brucei* [52].

To further investigate the enrichment pattern of G4 structures in the *T. brucei* genome, we quantified ChIP-Seq peak enrichment across annotated genomic features using a randomisation approach [21]. Specifically, we used *bedtools shuffle* to generate three independent sets of randomised peaks across the genome, aiming to preserve the number and length of original peaks but randomising their locations. By comparing the number of ChIP-Seq G4-peaks to random occurrence, we calculated fold-enrichment values for each genomic feature. Our analysis revealed the highest enrichment within coding regions, with approximately a 2-fold increase (1.90 ± 0.03), highlighting a potential functional role for G4 structures in regulating or interacting with protein-coding sequences in the *T. brucei* genome. The next most enriched features were TSS and TTS, exhibiting mean fold enrichments of 1.15 ± 0.03 and 1.13 ± 0.04, respectively. In contrast, intergenic regions, pseudogenes, and RNA genes showed little to no enrichment (fold enrichments of 0.05 ± 0.001, 0.15 ± 0.003, and 0.22 ± 0.08, respectively), suggesting these regions are less associated with detectable G4 formation.

These results suggest that biologically functional G4s in *T. brucei* might be primarily located within CDSs, as well as in TSS and TTS, with lower prevalence in intergenic regions and other genomic categories. Consistent with observations in the human genome, our data highlight the importance of experimental validation of G4 distribution through ChIP-Seq combined with comprehensive genomic annotation to accurately define the G4 landscape of pathogenic organisms with complex or repetitive genome architectures, such as *T. brucei*.

### *T. brucei* chromosome cores are abundant in G4 structures

Using our newly generated G4 map, we next aimed to evaluate the potential biological functions of the identified G4s. To achieve this, the relationship between G4 distribution and chromosome size in *T. brucei* was initially investigated using the set of 7,564 shared G4 peaks identified from ChIP-Seq data across three replicates. A radial plot revealed that the megabase chromosome core regions are densely packed with G4 peaks (Figure 3A). Additionally, chromosome size was found to be proportional to the number of peaks found, with a high R-squared value of 0.859 (Supplementary Figure S1, A). Considering that core regions of the megabase chromosomes are known to contain a higher number of gene clusters in comparison to subtelomeric regions, these findings suggest G4s may be relevant in *T. brucei* gene regulation. To rule out the possibility that the high frequency of G4s in the chromosome cores was due to differences in chromosome size or GC content, we normalised the number of G4 peaks for these two parameters. Specifically, the chromosome size (as number of bases) and GC content (as a percentage) were extracted from the FASTA file of the *T. brucei* reference genome using the ‘*awk’* command. Briefly, to normalise the data to the chromosome size (Snorm), we assigned the largest chromosome (i.e., chromosome 11) a size of 100 and normalised the other chromosome sizes relative to it (Snorm = [Size of chromosome / Size of largest chromosome] × 100). Then, the number of G4-peaks was divided by the normalised Snorm. The resulting values represent the number of peaks per normalised size unit, reflecting how densely distributed the G4-peaks are on the chromosome relative to its size. A similar normalization was applied for GC content, where an arbitrary value of 100 was assigned to the chromosome with the highest GC content (chromosome 10 core, with GC = 48.02%). The number of G4 peaks was then normalised to this GC content. This analysis revealed that G4s are more enriched in chromosome cores compared to subtelomeric and other chromosomal regions (Supplementary Figure S1).

**Figure 3:**
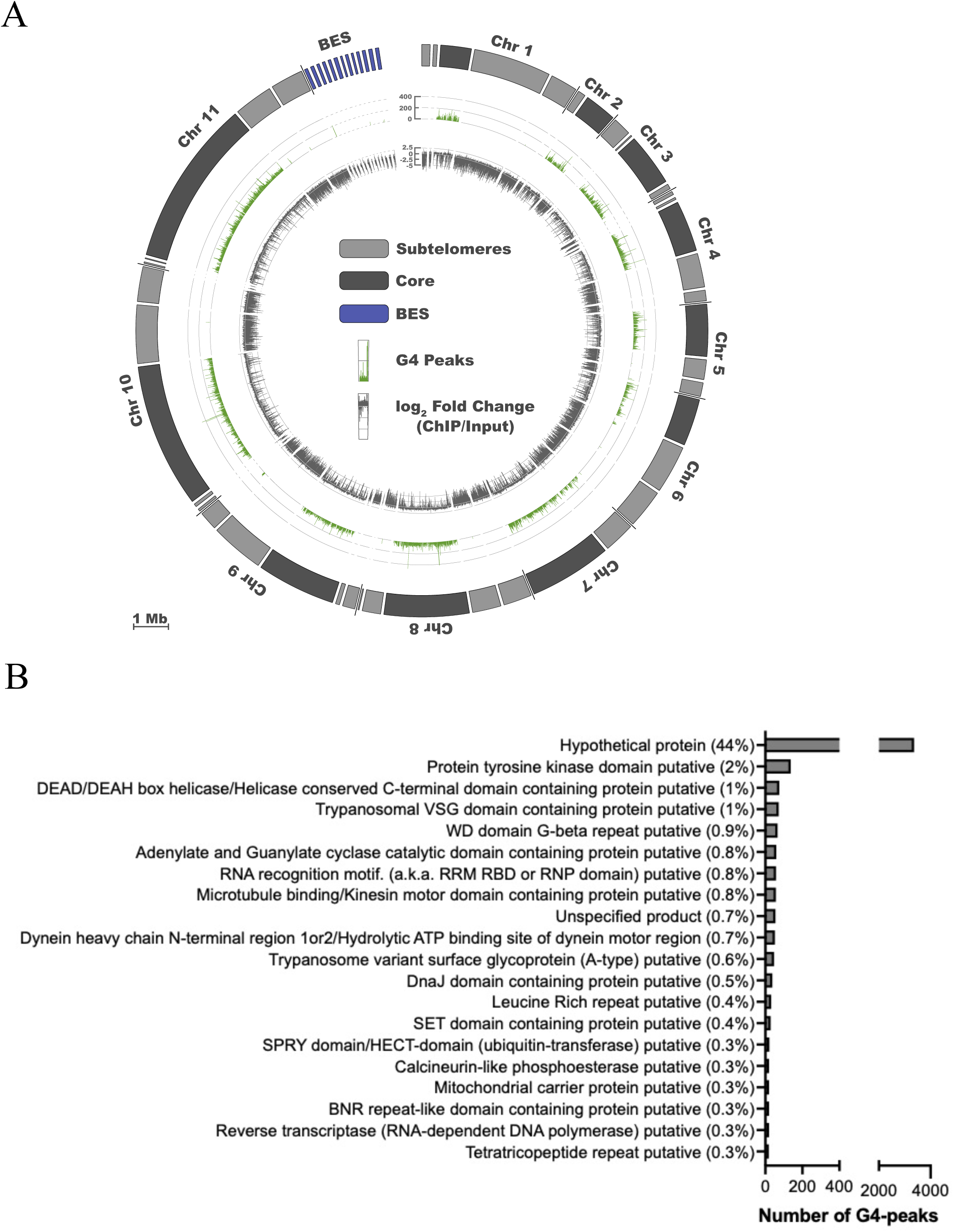
Genomic distribution and functional annotation of G4s in *T. brucei* chromosomes. A) Radial plot of ChIP-Seq data shows G4-enrichment across the *T. brucei* genome which consists of 11 megabase chromosomes and bloodstream expression sites (BESs). The inner track (grey) shows the ratio of ChIP/Input as log_2_ fold change across the genome, calculated using *BamCompare* with a bin size of 1000 bp. The outer track (green) represents the G4-peaks in each region, while peak height reflects the peak score as determined by *MACS2* analysis. B) Bar graph showing the most common functional categories associated with G4 peaks identified from ChIP-Seq data. The x-axis indicates the number of G4 peaks, and percentages represent the proportion of total peaks assigned to each category.

To infer functional roles of G4s in *T. brucei*, we analysed the most common gene functions associated with the 7,564 shared G4 peaks identified from ChIP-Seq data (Figure 3B). The largest proportion of these G4 peaks (44%) mapped to hypothetical proteins, indicating that a substantial portion of the G4-enriched regions in *T. brucei* are associated with genes of unknown function. In addition, we observed G4 peaks associated with protein tyrosine kinase domain (2%) and DEAD/DEAH box helicase-containing (1%) proteins, suggesting potential roles for trypanosomal G4s in gene expression regulation and RNA metabolism. Other enriched categories include trypanosomal VSG domain-containing (1%) and WD domain G-beta repeat (0.9%) proteins, which may reflect the involvement of G4s in antigenic variation, a key process in trypanosome immune evasion. This observation is of particular interest, as it could rationalise the reported high potency of G4 ligands in treating trypanosomiasis [41-49]. G4 peaks were also identified in genes encoding for adenylate and guanylate cyclase catalytic domain proteins (0.8%), RNA recognition motifs (0.8%), and microtubule-binding/kinesin motor proteins (0.8%). Less frequent associations were observed with dynein motor proteins, DnaJ domain-containing proteins, and leucine-rich repeat proteins (0.6– 0.4%). Interestingly, genes involved in RNA-dependent DNA polymerase activity (reverse transcriptase) and mitochondrial carrier proteins (0.3%) were also identified, suggesting a potential link between G4s and mitochondrial function or genome stability.

Our analysis highlighted the enrichment of G4 structures in critical pathways that may serve as potential targets for therapeutic intervention. To explore this further, we aimed to determine whether treatment with G4 ligands, some of which have been previously shown to inhibit parasite growth, could affect these pathways and justify the activity of these ligands as antiparasitic agents. Specifically, we sought to assess whether such treatment leads to changes in the expression profile of these genes, offering insights into how G4 stabilization might affect the growth and regulatory mechanisms of *T. brucei*.

### G4-ligands affect the viability of trypanosomes

To evaluate the effect of known G4-targeting ligands on *T. brucei* survival, we performed viability assays using two established G4-ligands, PhenDC3 [75] and CX-5461[30]. CX-5461 was initially developed by Hurley and co-workers as an anticancer agent targeting G4s. Subsequent studies revealed its additional activity as an RNA polymerase I (RNA Pol I) inhibitor [84]. Interestingly, CX-5461 has been shown to kill *T. brucei in vitro* and this activity has been previously linked to RNA Pol I inhibition rather than G4 stabilisation [40]. However, its potential to act as an antiparasitic agent based on its ability to target G4 structures has not been investigated yet. Thus, we hypothesised that CX-5461 might act both as a G4 binder and as a RNA Pol I inhibitor, while PhenDC3 would provide a more G4-driven mechanistic view on the antiparasitic potential linked to G4 targeting ligands.

We confirmed that *T. brucei* is sensitive to G4s ligands. CX-5461 exhibited the most potent antiparasitic activity with an EC_50_ of 0.15 ± 0.01 μM (Table 2). However, it also exhibited significant cytotoxicity in human HeLa cells (CC_50_ <0.20 μM), resulting in a low selectivity index (S.I.) of below 1.3, which indicates poor selectivity for *T. brucei* (Table 2). The low selectivity shown by CX-5461 may be due to its dual mechanism of action, whereby it also inhibits RNA Pol I, leading to potential off-target effects and reduced selectivity in human cells. In contrast, PhenDC3, showed potent killing activity against the parasites with a favourable selectivity profile, yielding an improved S.I. of 150 (*T. brucei* EC_50_: 0.36 ± 0.06 μM; HeLa CC_50_: 53 ± 3 μM). This result indicated that the targeting of G4 structures may translate into selective *T. brucei* killing, making PhenDC3 a promising molecular scaffold for potential drug development.

**Table 2.** Antiparasitic activity and cytotoxicity in HeLa cells of PDS and PhenDC3 (pure G4-ligands), CX-5461 (dual G4-ligand/RNA Pol I inhibitor), and BMH-21 (pure RNA Pol I inhibitor).

| Compound | *T. brucei*<br>EC<sub>50</sub> (μM) | *T. brucei*<br>EC<sub>90</sub> (μM) | HeLa<br>CC<sub>50</sub> (μM) | S.I.<br>HeLa/Tb<br>EC<sub>50</sub> |
| --- | --- | --- | --- | --- |
| **PhenDC3** | 0.36 ± 0.06 | 1.04 ± 0.06 | 53 ± 3 | 150 |
| **CX-5461** | 0.15 ± 0.01 | 0.19 ± 0.01 | <0.20* | <1.3 |
| **BMH-21** | 0.11 ± 0.01 | 0.17 ± 0.06 | <0.28* | <2.5 |

Cell viability of *T. brucei* and HeLa cells was assessed using Alamar Blue (resazurin) as described (Materials and Methods). EC<sub>50</sub> and CC<sub>50</sub> values were calculated from 11-point concentrations and reported as the mean ± standard deviation (SD). Assays were performed in triplicate. Selectivity Indices (S.I.) were calculated as HeLa CC<sub>50</sub> over *T. brucei* EC<sub>50</sub> values. \*For HeLa cells, lowest dose tested.

Next, we aimed to determine whether the high selectivity displayed by PhenDC3 could be ascribed to changes in expression of genes containing G4s, as detected by ChIP-Seq. To investigate this, we performed RNA-seq and evaluated the transcriptional responses elicited by these ligands on the *T. brucei* transcriptome. Total RNA was extracted from *T. brucei* cells treated with either CX-5461 or PhenDC3 and compared to an untreated control. To disentangle between the transcriptional response mediated by CX-5461 as a G4-binder from RNA Pol I inhibition, a specific RNA Pol I inhibitor, BMH-21, was also included in our analysis [40].

### PhenDC3 induces a distinct transcriptome profile compared to CX-5461 and BMH-21

RNA-seq was performed on *T. brucei* bloodstream form parasites after 4 hours treatment with PhenDC3, CX-5461 and BMH-21 at concentrations of 2 × EC_50_. Multidimensional Scaling (MDS) analysis of RNA-seq replicates revealed clear clustering of samples according to treatment condition (control, CX-5461, PhenDC3, and BMH-21), indicating high reproducibility and data quality, as well as distinct transcriptome profile to each compound (Supplementary Figure S2), as shown by samples forming well-separated clusters corresponding to their respective groups. Interestingly, a clear separation for the PhenDC3-treated samples from both the control and the other two treatments was observed, which indicates a difference in gene expression profile. This observation was confirmed by differential gene expression (DGE) analysis which revealed a distinct gene expression pattern for PhenDC3 compared to CX-5461 and BMH-2, as shown by the volcano plots (Figure 5A). DGE analysis was performed using a stringent false discovery rate (FDR) threshold of 0.05 and a log_2_ Fold Change (FC) of ± 1 (equivalent to a 2-fold change). The FC threshold allowed the identification of a broad set of potentially relevant genes for downstream analysis, while the FDR cut-off of 0.05 provides rigorous control for multiple testing, minimising the likelihood of false positives. Figure 4B and C summarise the findings from the DGE analysis, which included all gene types (i.e., protein coding genes, pseudogenes, and ncRNAs). Overall, irrespective of the treatment, approximately 98% (∼16,900 genes) of the total genes were not found to be differentially expressed compared to the untreated control. Among the differentially expressed genes, CX-5461 appeared to impact the highest number of genes (357), followed by PhenDC3 (294) and BMH-21 (289). Notably, PhenDC3 induced a nearly equal number of upregulated (139) and downregulated (155) genes, while CX-5461 and BMH-21 had a higher prevalence of downregulated genes, an effect that may be due to their inhibitory activity on the RNA Pol I function. Given the G4-targeting potential of PhenDC3, we investigated whether the involvement of G4-structures could explain the different expression profiles observed for this compound. To assess this, we intersected RNA-seq data with G4 ChIP-Seq data. As shown in Figure 4B, PhenDC3 resulted in a higher overall number of G4-containing genes being differentially expressed (43 genes total, of which 33 upregulated and 10 downregulated), compared to BMH-21 (16 total) or CX-5461 (15 total). Furthermore, PhenDC3 induced the strongest gene upregulation overall, including among G4-containing genes (33 genes), suggesting a potential role for G4-associated regulation of gene expression. CX-5461 and BMH-21 triggered broader gene downregulation, with a low proportion of G4-containing genes among downregulated genes, suggesting lower selectivity in targeting of G4s. Furthermore, none of the fifteen genes impacted by all three compounds harboured G4s. We also investigated the genes differentially expressed in response to both PhenDC3 and CX-5461, given their shared potential for G4 targeting. Among the 35 genes affected by both compounds, only one gene (*Tb427_110072300*) that contained a G4 motif and was upregulated in both treatments. This overlap, despite being limited to one gene, might reflect that CX-5461 can also act as a G4-ligand for certain targets. However, its global gene expression profile resembles more closely BMH-21, likely due to their shared mechanism of RNA Pol I inhibition. Given the distinct gene upregulation of G4-containing genes observed with PhenDC3, we examined the functional annotation of the 33 upregulated genes harbouring G4 motifs. While a large proportion of *T. brucei* genes are annotated as hypothetical proteins due to the lack of identifiable homologs in sequence databases, the methyltransferase domain annotation was unique to six G4-containing genes upregulated by PhenDC3. This domain was not found in any downregulated genes, nor in those up- or downregulated by CX-5461 or BMH-21. This points to a potential role of G4 structures in the regulation of methylation-related genes, something that has been previously reported in other eukaryotic organisms [85-87] and may suggest a regulatory mechanism involving epigenetic modulation mediated by G4s and perturbed upon PhenDC3 treatment.

**Figure 4:**
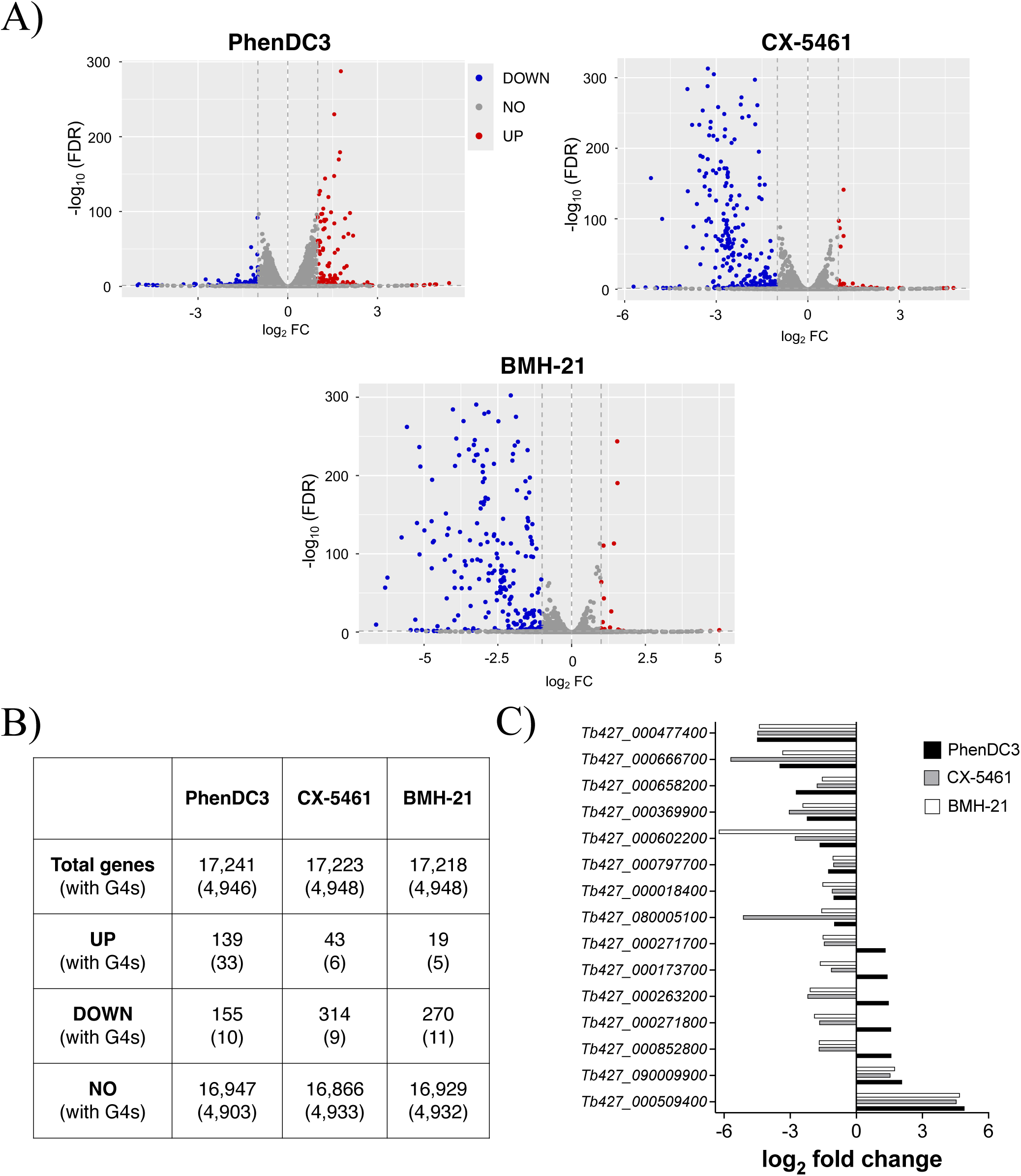
PhenDC3 induces a distinctive transcriptome profile in *T. brucei*. A) Volcano plots depicting differential gene expression profiles for PhenDC3, CX-5461, and BMH-21 treatments relative to untreated control (FDR < 0.05, log_2_FC ≥ 1). B) Table summarising the total number of genes (including protein coding genes, pseudogenes, and ncRNAs) and the count of differentially expressed genes (DEGs) for each treatment condition using an FDR threshold of 0.05 and a log_2_ fold change cut-off of ±1. UP = upregulated; DOWN = downregulated; NO = no significant change compared to untreated control. Number of genes carrying G4s are indicated in parentheses. C) Bar plot with the fifteen shared genes up- or down-regulated by PhenDC3, CX-5461 and BMH-21. Their expression levels are indicated as log2 fold change compared to untreated control.

**Figure 5:**
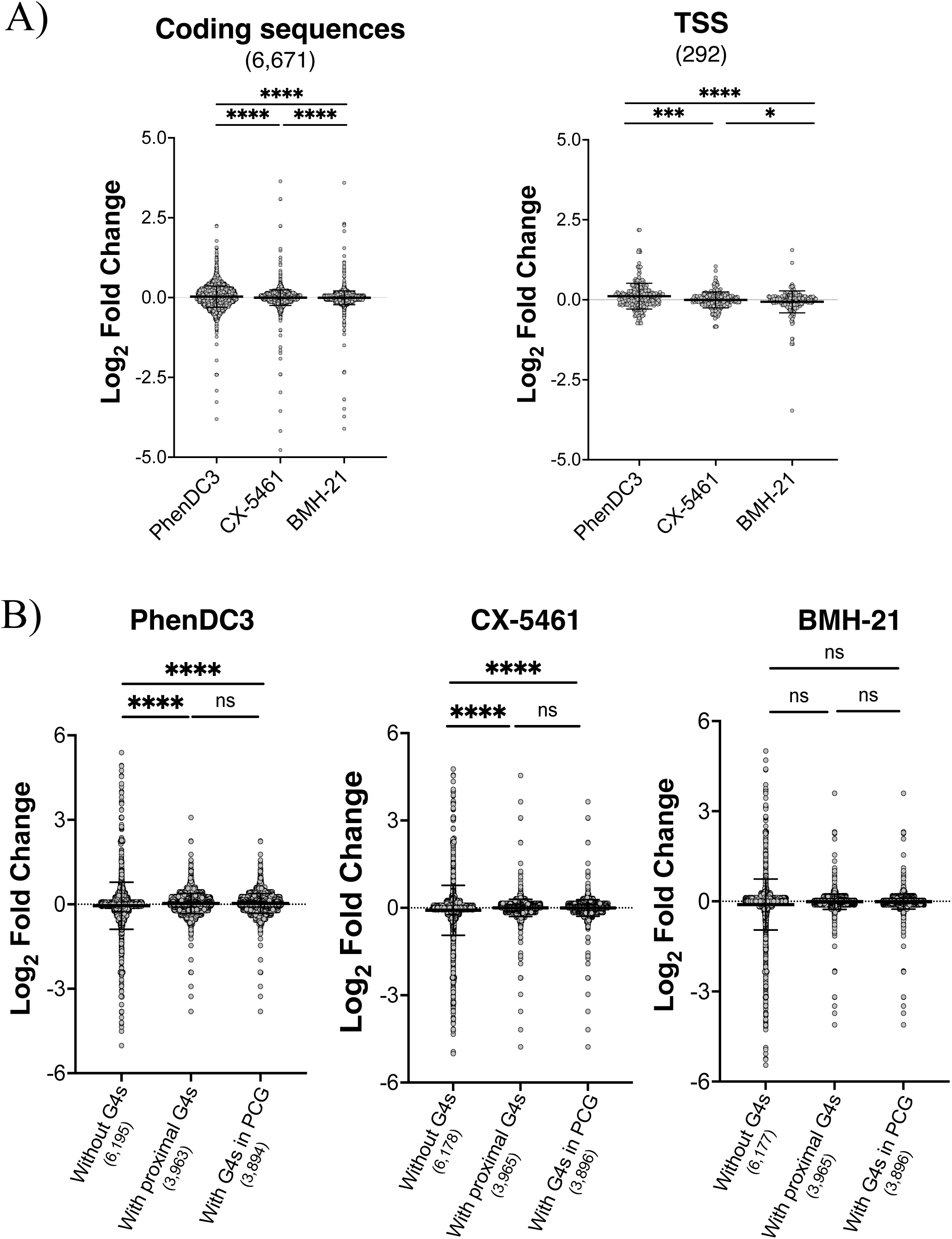
Expression changes of G4-associated genes across compound treatments. A) Gene expression changes of G4-containing genes, as identified by G4 ChIP-Seq, upon treatment with PhenDC3, CX-5461, or BMH-21 in coding sequences, transcriptional start site (TSS). The number of G4-associated genes per category is shown in parentheses. B) Global changes in gene expression of *T. brucei* genes with and without G4s upon compound treatment. Change in gene expression (as log2 fold change) of genes in coding sequences (protein coding genes, PCG). For both A) and B), differential gene expression relative to untreated control is shown as mean log2 fold change (FC) ± standard deviation (SD) for each treatment. Statistical significance between groups was determined using the Mann–Whitney U-test, where **** denotes p ≤ 0.0001, *** denotes p ≤ 0.001, ** denotes p ≤ 0.01, * denotes p ≤ 0.05, and ns indicates no significant difference (i.e., p > 0.05).

### G4-feature analysis revealed upregulation of G4-containing coding sequences to PhenDC3

Our DGE analysis highlighted a distinct gene expression pattern induced by the G4-ligand PhenDC3, compared to the RNA Pol I inhibitors CX5461 and BMH-21. Furthermore, a higher association between transcriptional dysregulation of genes harbouring G4s was observed for PhenDC3, suggesting a more specific G4-targeting mechanism.

To further evaluate the impact of each treatment on G4-associated genes, we specifically examined differences in gene expression induced by the three compounds on genes identified as carrying G4 motifs based on G4 ChIP-Seq data. Genes containing G4s were therefore analysed according to their genomic location, grouping them into: CDSs, CDSs overlapping either a TSS or TTS, pseudogenes, and non-coding RNAs (ncRNAs). As shown in Figure 5A, this analysis revealed that the highest differences in gene expression of G4-associated genes between compounds were observed particularly in CDSs and TSS regions, while a lower significance was observed in TTS, pseudogenes, and ncRNAs (Supplementary Figure S3). This finding is in agreement with our ChIP-Seq data showing an enrichment of G4s particularly in CDSs and TTS amongst the other categories, which may reflect that PhenDC3 has a strongest effect where G4s are more prevalent.

To further investigate this hypothesis, we assessed the influence on global gene expression induced by each compound by stratifying genes that do and do not contain G4s. Specifically, we annotated the RNA-Seq genes using the same genomic features defined for G4 ChIP-Seq analysis (see Material and Methods), categorising them into: CDSs, CDSs overlapping either a TSS or TTS, pseudogenes, and ncRNA. We then grouped the genes into three main categories: genes that do not contain G4s (*without G4s*), genes that contain or have proximal G4s (*with proximal G4s*, e.g., G4s in a near TSS/TTS), and genes that carry a G4 (*with G4s*). Figure 5B shows fold changes in gene expression for genes associated in regions with and without G4s upon exposure to each treatment. Interestingly, we observed that PhenDC3 and CX-5461 exhibit a significant difference in expression between gene with and without G4s particularly in coding sequences (Figure 5B). In contrast, BMH-21, does not show a significant difference in expression between genes with and without G4s in coding sequences (Figure 5B). Interestingly, while CX-5461 and BMH-21 induced an overall downregulation of genes (both without and with G4s), PhenDC3 induced an opposite effect, downregulating genes without G4s and upregulating genes carrying G4s. Notably, this effect induced by PhenDC3 was particularly significant in CDS regions, where G4s were found to be mostly enriched, while it was found to be less significant in other genomic locations, such as TSS, TTS, pseudogenes, and ncRNA (Supplementary Figure S4). This evidence further supports our hypothesis that G4s are enriched at CDSs in *T. Brucei* and that treatment with G4-ligands, such as PhenDC3, may act through a G4-mediated mechanism that alters preferentially genes carrying G4s. We also observed a consistent trend in the expression changes produced by PhenDC3 in the three categories, with the highest G4-enrichment as determined by G4 ChIP-Seq data (i.e., CDS, TSS, and TTS regions) corresponding to a significant upregulation of genes carrying G4s (Figure 5B and Supplementary Figure S4), while producing an overall downregulation in genomic locations where G4s are less prevalent (pseudogenes and ncRNAs, Supplementary Figure S4). In contrast, CX-5461 and BMH-21 showed a more variable effect on gene expression across the different genomic locations examined, which cannot be ascribed to G4-targeting (Figure 5B and Supplementary Figure S4).

## DISCUSSION

Our study presents the first comprehensive analysis of G-quadruplex (G4) DNA structures in trypanosomatid parasites, providing insights into G4’s potential as molecular targets to treat infections caused by these parasites.

Through whole-genome *in silico* analyses, we identified putative G4-quadruplex sequences (PQSs) across the genomes of 64 trypanosomatid species, revealing intriguing patterns of G4 richness that may reflect different biological functions mediated by G4s in different trypanosomatid species (Figure 1). For instance, the 23 *Leishmania* species analysed exhibited a consistently high PQS frequency (with an average of 3.74), and a narrow frequency range of 2.52 – 4.57, suggesting that G4s might have conserved functional roles. Conversely, the 23 *Trypanosoma* species examined showed a broader range of PQS frequencies ranging between 0.85 – 4.34 PQS/Kb. We observed a near-linear relationship between GC content and PQS frequency across genera, supporting the well-established notion that G4s are more likely to form in GC-rich regions [21]. However, we identified a notable exception to this trend, which suggested that GC content alone does not fully explain the prevalence of predicted G4 structures. *Porcisia* species retained the largest proportion of PQSs at increasing G4Hunter thresholds, despite having a lower GC content compared to other related species, such as *Leishmania* (Figure 1C). This suggests that other factors affect G4 prevalence in these organisms that go beyond GC richness. For example, *Leishmania* is a medically significant genus that infects both humans and animals, relying on a broad repertoire of cell surface proteins to facilitate the intracellular propagation of amastigotes within vertebrate macrophages—a critical adaptation for their survival and virulence [88]. In contrast, *Porcisia* species, which infect porcupines, have a substantially reduced repertoire of such proteins and do not rely on macrophage infection for their lifecycle [88]. The retention of PQSs in *Porcisia* despite lower GC content might reflect alternative mechanisms regulated by G4s that are tailored around the distinct ecological niche of these species.

To gain further insights on the potential physiological relevance of G4s in these organisms, we next applied G4 ChIP-Seq to map G4s in *T. brucei* chromatin, generating the first chromatin-based map of G4s in this parasite. G4 ChIP identified 7,564 G4-peaks consistent across 3 distinct replicates (Figure 2C), confirming the particularly dense distribution of G4 structures in *T. brucei* already predicted computationally. This is particularly relevant when considering the relatively small size of the *T. brucei* genome (26 Mb) compared to other species, such as humans (3.1 Gb). This further highlighted the potential for functional relevance of G4s in *T. brucei* and agrees with the established potential of G4 ligands in acting as antiparasitic agents. The genome-wide distribution of G4s detected by ChIP-Seq differs markedly from what has been observed in human, whereby G4s are predominantly observed in introns, intergenic, and promoter regions [21,52]. In contrast to previous findings, our data reveal a striking enrichment of *T. brucei* G4s within coding sequences, with comparatively lower abundance at TSS, TTS and other regions (Figure 2D and E). This distribution diverges from that reported by Marsico *et al.* using G4-seq, which showed a pattern more closely resembling the G4 landscape in human genomes when measured on naked single-stranded DNA (G4-Seq). These differences likely reflect the distinct experimental approaches employed. G4-seq is based on sequencing extracted genomic DNA under conditions that promote G4 formation (e.g., potassium ions and/or G4-stabilising ligands) and identifies G4s by detecting mismatches between sequencing reads generated under these conditions. Our ChIP-based approach, in contrast, captures G4s within native chromatin extracted directly from *T. brucei* parasites. As such, it may better reflect the physiological and dynamic nature of G4 formation *in vivo*. Interestingly, despite these methodological differences, the total number of G4s identified in our study (7,564 peaks) is broadly comparable to the range reported by Marsico *et al.* [21]. However, the genomic distribution of G4s in our ChIP-Seq data differs markedly from that observed in the G4-seq data., which may also result from differences in annotation strategies. While Marsico *et al.* applied a cross-species framework, our analysis was tailored to the unique genomic architecture of *T. brucei*, including its organisation into polycistronic transcription units.

Altogether, these observations underscore the influence of experimental context— particularly the chromatin state and genome annotation—on G4 detection. G4-seq, which uses naked single-stranded DNA, does not account for chromatin structure or other dynamic factors that may regulate G4 folding and stability in a cellular context. By contrast, ChIP-seq approaches preferentially capture G4s that are stably folded under physiological conditions, providing a potentially more biologically relevant snapshot of G4 dynamics in living cells.

The high density of G4s in coding sequences of chromosome cores of the *T. brucei* genome suggests that these structures might be involved in the regulation of key biological functions of the parasite, including transcriptional regulation of genes. Therefore, we investigated the effects of established G4-ligands on the transcriptional activity of the parasite. The sensitivity of *T. brucei* to G4-ligands highlights the therapeutic potential of these compounds in treating trypanosome infections, as also evidenced by the recent reports of various G4-ligands with anti-trypanosomal activity [40-50] (Table 2). PhenDC3, exhibited potent antiparasitic activity with a favourable selectivity profile, making it a strong candidate for further development of antiparasitic drugs. In contrast, the dual G4-ligand and RNA Pol I inhibitor CX-5461 showed high anti-trypanosomal efficacy but low selectivity, which is in agreement with its use in humans as an anticancer agent. The distinct transcriptome profiles following PhenDC3, CX-5461, and BMH-21 exposure revealed by RNA-seq further supports their different modes of action. The ability of PhenDC3 to induce changes in the expression of G4-containing genes suggests a relevant role of G4s in regulating gene expression levels in *T. brucei* (Figure 4 and 5), which is in agreement with what has been extensively reported in human cells [89-91]. The significant enrichment of DGEs in coding sequences containing G4s, particularly upon PhenDC3 and CX-5461 treatment, further supports the hypothesis that G4s have a role as modulators of gene expression in trypanosomes (Figure 5B). Targeting G4 structures offers a promising strategy for perturbing gene expression homeostasis in pathogens like *T. brucei*, which can be potentially leveraged to overcome the increasing incidence of resistance against current therapeutic agents. We anticipate that this approach could also have broader applications in combating infectious diseases beyond parasites and might be explored in broader microbiology contexts.

The distinct upregulation of G4-containing genes observed following PhenDC3 treatment might be explained by the different genomic distribution of G4s in *T. brucei* compared to mammals. Additionally, this could also reflect different mechanisms of G4-protein recognition leveraged in *T. brucei* that might be enhanced in the presence of the ligand, leading to gene expression stimulation. This would be in line with our recent finding demonstrating that G4 ligands can act as molecular glues and increase G4-protein interactions at certain G4-sites [92]. From a more functional perspective, the upregulation of methyltransferase domain-containing proteins, which is unique to PhenDC3 and absent for CX-5461 or BMH-21, warrants further investigation and highlights potential for epigenetic relevance in parasite biology. Methyltransferase-associated G4 peaks constitute a modest fraction of total G4 ChIP-Seq peaks (13 identified) and exhibit high peak scores, suggesting stable G4 formation at these *loci*. Furthermore, RNA-seq analysis identified 10 methyltransferase genes harbouring G4 motifs that were specifically upregulated by PhenDC3 but remained largely unaffected by the other compounds. This confirms a link between G4 stabilization and epigenetic regulation mediated by methyltransferase activity, which has been previously described in other organisms [86,93].

Overall, this study provided the first chromatin-based genome-wide map of G4s in *T. brucei*, highlighting species-specific distribution, transcriptional relevance, and therapeutic potential of G4s in trypanosomes. The distinct G4 enrichment pattern observed in *T. brucei*, compared to computation predictions and other mapping methods, reinforces the importance of experimental validation using physiologically relevant approaches when evaluating G4 prevalence—an observation previously made in human cells. Future studies should now focus on exploring the mechanistic roles of unknown proteins that have shown potential for gene expression modulation by means of G4-targeting, as this might lead to the rational design of a new generation of therapeutic agents. Additionally, further optimisation of current G4-ligands with improved selectivity profiles holds promise for the development of new targeted therapies against trypanosome infections.

## ACKNOWLEDGEMENTS

We acknowledge Prof. Gloria Rudenko and her research group at Imperial College London for invaluable guidance through the initial phases of the study.

## SUPPLEMENTARY DATA

Supplementary Data are available online.

