## Supplementary Information for "Genome-wide mapping of DNA G-quadruplexes in *Trypanosoma brucei* chromatin reveals enrichment in coding regions"

**SUPPLEMENTARY DATA**

**Table S1.** G4Hunter analysis of putative G-quadruplex sequences (PQSs) identified in the genomes of trypanosomatids and control species (Schistosoma and Homo sapiens).

**Table S2.** Comparison of predicted and experimental G4-forming sequences in T. brucei chromosomes.

**Table S3.** Functional analysis of differentially expressed genes in T. brucei

**Figure S1.** Normalised G4 peaks across T. brucei chromosomes.

**Figure S2.** Multidimensional scaling (MDS) plot of RNA-seq data.

**Figure S3.** Gene expression changes of G4-containing genes upon treatment with PhenDC3, CX-5461, or BMH-21.

**Figure S4.** Global changes in gene expression of T. brucei genes with and without G4s upon compound treatment.

**Table S1. G4Hunter[1-3] analysis of putative G-quadruplex sequences (PQSs) identified in the genomes of trypanosomatids and control species (*Schistosoma* and *Homo sapiens*).** A threshold of 1.2 and a minimum sequence length of 25 nucleotides were applied in this analysis. Genomic details, including genome length (in base pairs), GC content (percentage), and accession numbers, are provided. [a]As reported in Supp. ref.[1]; [b]as reported in Supp. ref. [2]

| Genus | Species | Genome size (bp) | Number of PQS | Frequency of PQSs per 1000 base pairs | GC Content (%) | Accession Number |
|---|---|---|---|---|---|---|
| *Crithidia* | *C. thermophilia* | 29,959,113 | 53,466 | 1.785 | 54.07 | GCA_030849055.1 |
| | *C. brevicula* | 35,360,925 | 66,534 | 1.882 | 53.42 | GCA_030849845.1 |
| | *C. sp. LVH-60A* | 34,393,440 | 92,369 | 2.686 | 57.23 | GCA_030078075.1 |
| | *C. fasciculata* | 41,297,378 | 104,539 | 2.531 | 57.01 | GCA_000331325.2 |
| | *C. mellificae* | 49,951,730 | 94,942 | 1.901 | 53.51 | GCA_002216565.1 |
| | *C. acanthocephali* | 33783172 | 62,084 | 1.838 | 56.22 | GCA_000482105.1 |
| | *C. expoeki* | 34,077,985 | 61,301 | 1.799 | 54.40 | GCA_900240875.1 |
| *Herpetomonas* | *H. samuelpessoai* | 32,248,164 | 91,995 | 2.853 | 56.37 | GCA_030849835.1 |

| | | | | | | |
|---|---|---|---|---|---|---|
| | *H. muscarum* | 30,844,810 | 156,425 | 5.071 | 58.20 | GCA_000482205.1 |
| | *H. tarakana* | 26,680,300 | 150,158 | 5.628 | 61.21 | GCA_030849825.1 |
| *Homo* (human) | *H. sapiens* | 3,095,690,000[a] | 2,890,423[a] | 0.93 | 37.8[a] | - |
| *Leishmania* | *L. major* | 32,855,089 | 123,264 | 3.752 | 59.72 | GCA_000002725.2 |
| | *L. brazilienis* | 32,068,771 | 120,841 | 3.768 | 57.62 | GCA_000002845.2 |
| | *L. infantum* | 32,122,061 | 116,863 | 3.638 | 59.54 | GCA_000002875.2 |
| | *L. donovani* | 32,444,968 | 113,086 | 3.485 | 57.31 | GCA_000227135.2 |
| | *L. mexicana* | 32,108,741 | 128,660 | 4.007 | 59.74 | GCA_000234665.4 |
| | *L. panamensis* | 30,688,794 | 112,805 | 3.676 | 56.25 | GCA_000755165.1 |
| | *L. aethiopica* | 33,648,436 | 132,773 | 3.946 | 60.38 | GCA_003992445.1 |
| | *L. amazonesis* | 32,156,470 | 126,536 | 3.935 | 59.50 | GCA_005317125.1 |
| | *L. guyanensis* | 32,544,702 | 127,342 | 3.913 | 57.88 | GCA_024970365.1 |
| | *L. lainsoni* | 34,152,029 | 129,606 | 3.795 | 57.84 | GCA_003664395.1 |

| | | | | | | |
|---|---|---|---|---|---|---|
| | *L. tropica* | 32,700,668 | 121,613 | 3.719 | 59.67 | GCA_014139745.1 |
| | *L. chagasi* | 31,924,566 | 116,007 | 3.634 | 59.54 | GCA_014466975.1 |
| | *L. martiniquensis* | 32,413,670 | 148,059 | 4.568 | 59.85 | GCA_017916325.1 |
| | *L. orientalis* | 34,194,276 | 131,793 | 3.854 | 59.72 | GCA_017916335.1 |
| | *L. naiffi* | 31,671,391 | 120,914 | 3.818 | 57.21 | GCA_962239345.1 |
| | *L. lindenbergi* | 31,126,267 | 117,855 | 3.786 | 57.37 | GCA_962240435.1 |
| | *L. utingensis* | 31,161,167 | 119,308 | 3.829 | 57.38 | GCA_962240445.1 |
| | *L. shawi* | 31,105,824 | 117,500 | 3.777 | 57.31 | GCA_962240455.1 |
| | *L. sp. AIIMS/LM/SS/PKDL/LD-974* | 27,848,322 | 70,192 | 2.521 | 55.79 | GCA_000981925.2 |
| | *L. enriettii* | 33,318,864 | 123,067 | 3.694 | 59.57 | GCA_017916305.1 |
| | *L. sp. Ghana* | 35,953,538 | 141,312 | 3.930 | 59.67 | GCA_017918215.1 |
| | *L. sp. namibia* | 34,118,624 | 131,918 | 3.866 | 59.53 | GCA_017918225.1 |

| | | | | | | |
|---|---|---|---|---|---|---|
| | *L. tarentolae* | 31,932,047 | 101,820 | 3.189 | 57.03 | GCA_033953505.1 |
| *Leptomonas* | *L. pyrrhocoris* | 30,379,903 | 71,638 | 2.358 | 56.44 | GCA_001293395.1 |
| | *L. seymouri* | 27,764,161 | 70,625 | 2.544 | 55.39 | GCA_001299535.1 |
| *Phytomonas* | *P. sp. isolate EM1* | 17,780,869 | 38,130 | 2.144 | 43.46 | GCA_000582765.1 |
| | *P. serpens* | 25,693,183 | 66,683 | 2.595 | 46.56 | GCA_000331125.1 |
| | *P. francai* | 17,721,985 | 44,953 | 2.537 | 47.66 | GCA_001766655.1 |
| | *P. sp. isolate Hart1* | 18,129,152 | 38,448 | 2.121 | 41.12 | GCA_000982615.1 |
| *Porcisia* | *P. hertigi* | 34,958,538 | 152,816 | 4.371 | 56.02 | GCA_017918235.1 |
| | *P. deanei* | 29,506,589 | 122,855 | 4.164 | 55.10 | GCA_018683835.1 |
| *Schistosoma* | *S. mansoni* | 409,579,008 | 88,178[b] | 0.22 | 34.66[b] | - |
| | *S. japonicum* | 402,743,189 | 84,683[b] | 0.21 | 31.23[b] | - |
| *Trypanosoma* | *T. cruzi* | 89,937,456 | 176,243 | 1.960 | 51.55 | GCA_000209065.1 |
| | *T. grayi* | 20,934,132 | 45,745 | 2.185 | 53.59 | GCA_000691245.1 |
| | *T. theileri* | 29,822,211 | 25,307 | 0.849 | 34.52 | GCA_002087225.1 |

| | | | | | | |
|---|---|---|---|---|---|---|
| | *T. congolense* | 41,233,446 | 88,271 | 2.141 | 47.42 | GCA_002287245.1 |
| | *T. rangeli* | 21,157,315 | 54,990 | 2.599 | 51.52 | GCA_003719475.1 |
| | *T. conorhini* | 21,334,213 | 81,817 | 3.835 | 56.71 | GCA_003719485.1 |
| | *T. evansi* | 25,432,160 | 43,419 | 1.707 | 46.53 | GCA_917563935.1 |
| | *T. vivax* | 67,823,889 | 186,308 | 2.747 | 53.64 | GCA_021307395.1 |
| | *T. melophagium* | 23,303,718 | 27,578 | 1.183 | 41.23 | GCA_022059095.1 |
| | *T. brucei brucei 927* | 26,075,494 | 43,794 | 1.680 | 46.43 | GCA_000002445.1 |
| | *T. pestanai* | 30,657,805 | 52,320 | 1.707 | 43.23 | GCA_964197905.1 |
| | *T. boissoni* | 22,199,164 | 40,055 | 1.804 | 46.06 | GCA_030849725.1 |
| | *T. musculi* | 20,531,481 | 38,053 | 1.853 | 46.49 | GCA_036321165.1 |
| | *T. mega* | 27,412,799 | 53,162 | 1.939 | 44.16 | GCA_030849715.1 |
| | *T. avium* | 22,090,251 | 26,297 | 1.190 | 31.20 | GCA_030849755.1 |
| | *T. lewisi* | 20,726,896 | 38,309 | 1.848 | 46.47 | GCA_036321185.1 |
| | *T. platydactyli* | 20,548,520 | 45,378 | 2.208 | 48.81 | GCA_030849675.1 |

| | | | | | | |
|---|---|---|---|---|---|---|
| | *T. scelopori* | 20,296,703 | 58,309 | 2.873 | 52.09 | GCA_030849745.1 |
| | *T. erneyi* | 33,887,235 | 147,098 | 4.341 | 55.83 | GCA_964197915.1 |
| | *T. caninum* | 19,024,056 | 26,408 | 1.388 | 41.29 | GCA_036321205.1 |
| | *T. suis* | 57,600,987 | 153,130 | 2.658 | 48.35 | GCA_964197925.1 |
| | *T. brucei gambiense* | 22,148,088 | 40,944 | 1.849 | 47.09 | GCA_000210295.1 |

**Table S2. Comparison of predicted and experimental G4-Forming sequences in *T. brucei* chromosomes.** Overlap of G4-forming sequences in *Trypanosoma brucei* chromosomes: comparison of predictive analyses using *G4Hunter* [3-5] at thresholds of 1.2, 1.5, and 1.8 (with a 25-base nucleotide length) and experimental data from G4 ChIP-seq.

| **Chromosome** | **ChIP-seq n. of G4-peaks** | **G4Hunter 1.2** | **ChIP-seq ∩ G4Hunter** | **% Overlap** | **AVG % overlap** | **G4Hunter 1.5** | **ChIP-seq ∩ G4Hunter** | **% Overlap** | **AVG % overlap** | **G4Hunter 1.8** | **ChIP-seq ∩ G4Hunter** | **% Overlap** | **AVG % overlap** |
|---|---|---|---|---|---|---|---|---|---|---|---|---|---|
| **Chr11_core** | 1694 | 8595 | 924 | 54.5 | **36%** | 2606 | 322 | 19.0 | **11%** | 949 | 99 | 5.8 | **3%** |

| | | | | | | | | | | | | |
|---|---|---|---|---|---|---|---|---|---|---|---|---|
| **Chr10_core** | 1466 | 7739 | 828 | 56.5 | | 2327 | 291 | 19.8 | | 879 | 90 | 6.1 |
| **Chr8_core** | 849 | 4575 | 499 | 58.8 | | 1375 | 191 | 22.5 | | 485 | 65 | 7.7 |
| **Chr7_core** | 679 | 4053 | 407 | 59.9 | | 1249 | 162 | 23.9 | | 459 | 51 | 7.5 |
| **Chr9_core** | 632 | 3828 | 362 | 57.3 | | 1175 | 131 | 20.7 | | 465 | 37 | 5.9 |
| **Chr4_core** | 462 | 2557 | 242 | 52.4 | | 766 | 82 | 17.7 | | 279 | 24 | 5.2 |
| **Chr3_core** | 451 | 2669 | 290 | 64.3 | | 852 | 126 | 27.9 | | 278 | 39 | 8.6 |
| **Chr5_core** | 395 | 2464 | 226 | 57.2 | | 710 | 79 | 20.0 | | 305 | 25 | 6.3 |
| **Chr6_core** | 272 | 2100 | 145 | 53.3 | | 616 | 58 | 21.3 | | 225 | 21 | 7.7 |
| **Chr1_core** | 251 | 1591 | 150 | 59.8 | | 472 | 56 | 22.3 | | 205 | 15 | 6.0 |
| **Chr2_core** | 237 | 1517 | 137 | 57.8 | | 469 | 47 | 19.8 | | 185 | 11 | 4.6 |
| **Chr7_5A** | 106 | 435 | 23 | 21.7 | | 402 | 2 | 1.9 | | 26 | 0 | 0.0 |
| **Chr6_3A** | 13 | 556 | 1 | 7.7 | | 110 | 1 | 7.7 | | 37 | 0 | 0.0 |
| **Chr9_3B** | 11 | 347 | 4 | 36.4 | | 100 | 1 | 9.1 | | 21 | 0 | 0.0 |
| **Chr11_3B** | 5 | 549 | 3 | 60.0 | | 110 | 0 | 0.0 | | 19 | 0 | 0.0 |
| **BES1** | 2 | 50 | 1 | 50.0 | | 84 | 0 | 0.0 | | 13 | 0 | 0.0 |
| **BES15** | 2 | 30 | 0 | 0.0 | | 153 | 0 | 0.0 | | 2 | 0 | 0.0 |

| | | | | | | | | | | | | |
|---|---|---|---|---|---|---|---|---|---|---|---|---|
| **Chr10_3A** | 2 | 849 | 0 | 0.0 | | 450 | 0 | 0.0 | | 49 | 0 | 0.0 |
| **Chr11_3A** | 2 | 644 | 1 | 50.0 | | 311 | 1 | 50.0 | | 34 | 0 | 0.0 |
| **Chr5_3B** | 2 | 202 | 0 | 0.0 | | 197 | 0 | 0.0 | | 13 | 0 | 0.0 |
| **BES12** | 1 | 34 | 0 | 0.0 | | 164 | 0 | 0.0 | | 3 | 0 | 0.0 |
| **Chr1_3B** | 1 | 378 | 0 | 0.0 | | 66 | 0 | 0.0 | | 14 | 0 | 0.0 |
| **Chr3_3A** | 1 | 123 | 1 | 100.0 | | 15 | 0 | 0.0 | | 3 | 0 | 0.0 |
| **Chr5_3A** | 1 | 285 | 0 | 0.0 | | 199 | 0 | 0.0 | | 18 | 0 | 0.0 |
| **Chr6_3B** | 1 | 486 | 0 | 0.0 | | 173 | 0 | 0.0 | | 19 | 0 | 0.0 |
| **Chr8_3A** | 1 | 294 | 0 | 0.0 | | 139 | 0 | 0.0 | | 13 | 0 | 0.0 |
| **Chr8_5B** | 1 | 452 | 0 | 0.0 | | 184 | 0 | 0.0 | | 32 | 0 | 0.0 |

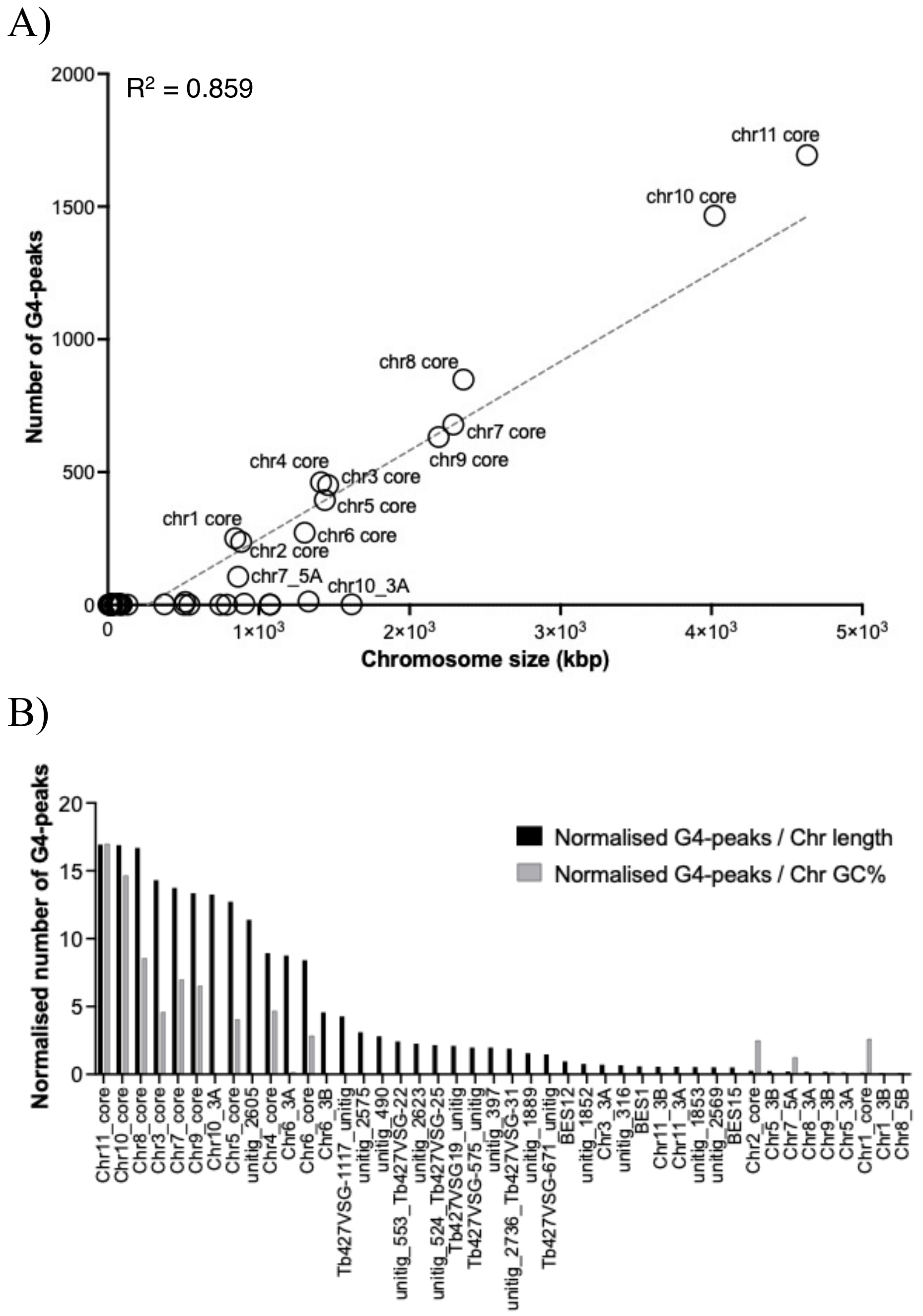

**Figure S1. Normalised G4 peaks across *T. brucei* chromosomes.** A) Logarithmic plot showing the number of G4 peaks identified from ChIP-Seq data as a function of chromosome size. A positive linear correlation indicates that the number of G4 peaks scales with increasing chromosome size. B) Black bars represent number of G4 peaks normalised to chromosome length (size), while grey bars are normalised to chromosome's GC content (as percentage). Despite normalisation, core chromosomes show a significant enrichment of G4 peaks compared to other regions.

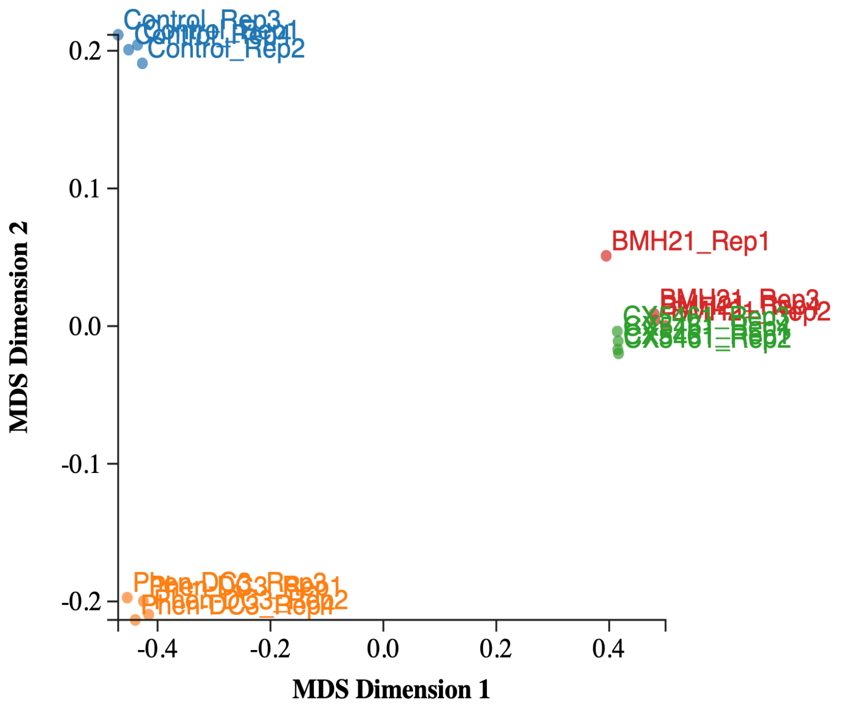

**Figure S2.** Multidimensional scaling (MDS) plot of RNA-seq data showing clustering of four technical replicates for *T. brucei* cells treated for 4 hours at 2 × $IC_{50}$ with either Phen-DC3 (orange), CX-5461 (green), BMH-21 (red), or untreated control (blue).

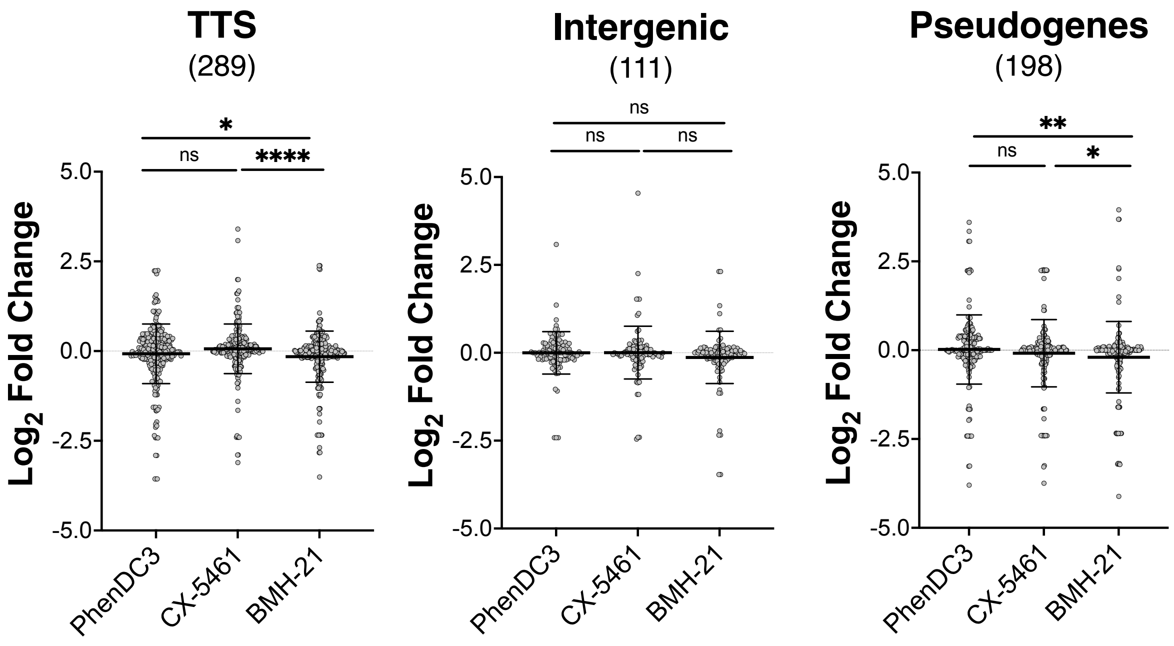

**Figure S3.** Gene expression changes of G4-containing genes, as identified by G4 ChIP-Seq, upon treatment with PhenDC3, CX-5461, or BMH-21 in transcriptional termination sites (TTS), intergenic regions and pseudogenes. The number of G4-associated genes per category is shown in parenthesis. Differential gene expression relative to untreated control is shown as mean $\log_2$ fold change (FC) ± standard deviation (SD) for each treatment. Statistical significance between groups was determined using the Mann–Whitney U-test, where **** denotes $p \leq 0.0001$, ** denotes $p \leq 0.01$, * denotes $p \leq 0.05$, and ns indicates no significant difference (i.e., $p > 0.05$).

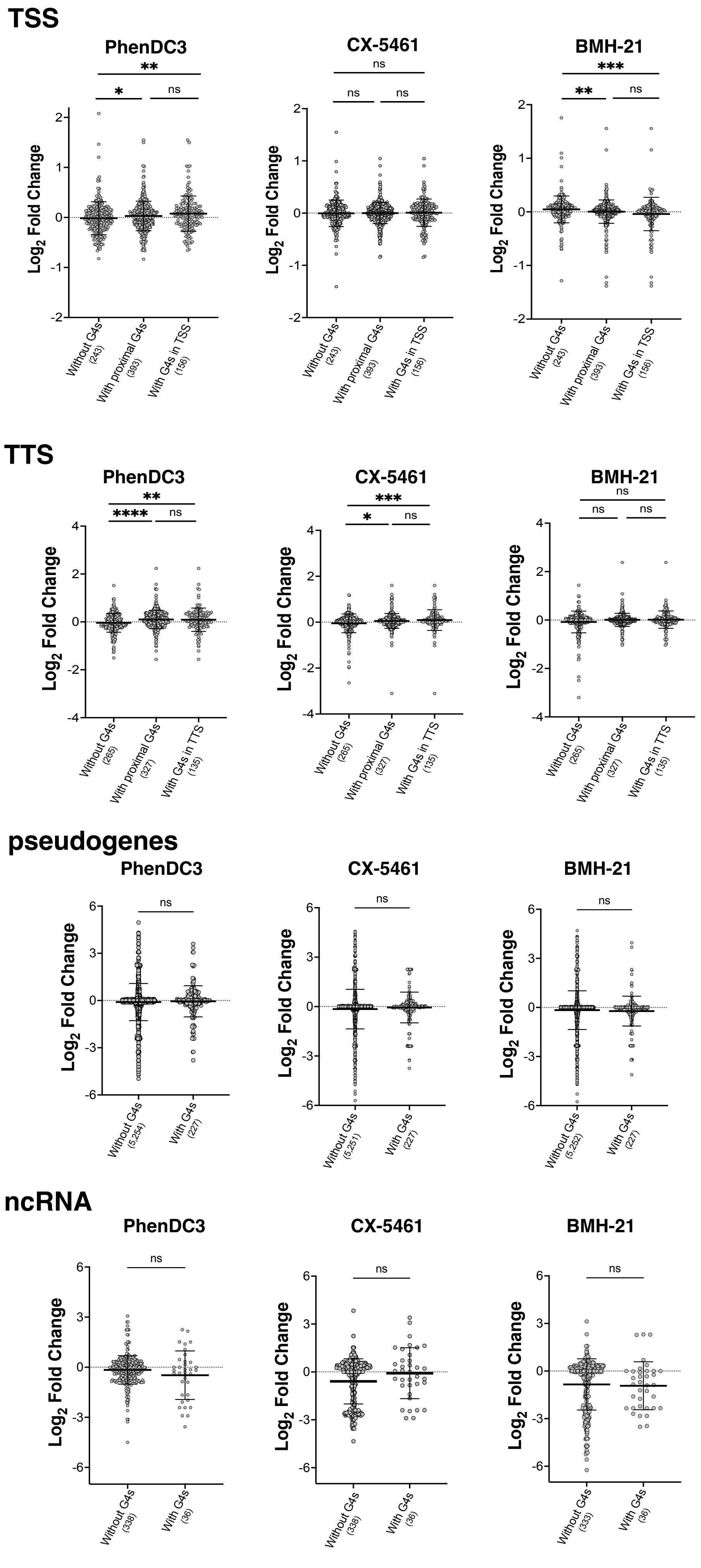

TSS
PhenDC3
CX-5461
BMH-21
Log2 Fold Change
Without G4s (243)
With proximal G4s (393)
With G4s in TSS (156)
TTS
Without G4s (265)
With proximal G4s (327)
With G4s in TTS (135)
pseudogenes
Without G4s (5,254)
Without G4s (5,251)
Without G4s (5,252)
With G4s (227)
ncRNA
Without G4s (338)
Without G4s (333)
With G4s (36)
ns
*
**
***
****

**Figure S4.** Global changes in gene expression of *T. brucei* genes with and without G4s upon compound treatment. Change in gene expression (as $\log_2$ fold change) of genes in transcriptional start sites (TSS), transcriptional termination sites (TTS), pseudogenes and non-coding RNAs (ncRNA). Differential gene expression relative to untreated control is shown as mean $\log_2$ fold change (FC) ± standard deviation (SD) for each treatment. Statistical significance between groups was determined using the Mann–Whitney U-test, where **** denotes $p \leq 0.0001$, *** denotes $p \leq 0.001$, ** denotes $p \leq 0.01$, * denotes $p \leq 0.05$, and ns indicates no significant difference (i.e., $p > 0.05$).
